# SERPINB3 induces the basal-like/squamous subtype and enhances disease progression in pancreatic cancer

**DOI:** 10.1101/2023.03.29.534766

**Authors:** Yuuki Ohara, Wei Tang, Huaitian Liu, Shouhui Yang, Tiffany H. Dorsey, Helen Cawley, Paloma Moreno, Azadeh Azizian, Jochen Gaedcke, B. Michael Ghadimi, Nader Hanna, Stefan Ambs, S. Perwez Hussain

**Author notes:** Address Correspondence to: S. Perwez Hussain, Ph.D., Chief, Pancreatic Cancer Section Laboratory of Human Carcinogenesis, Bldg. 37 Room 3044B, National Cancer Institute, NIH, 37 Convent Dr., Bethesda, MD 20892, Yuuki Ohara, M.D., Ph.D., Bldg. 37 Room 3054, National Cancer Institute, NIH, Wei Tang, Ph.D., Bldg. 37 Room 3050, National Cancer Institute, NIH.

## Abstract

Pancreatic cancer is a heterogeneous disease with distinct subtypes. Here, we investigated candidate driver genes of the highly aggressive basal-like/squamous molecular subtype of pancreatic ductal adenocarcinoma (PDAC). Integrative transcriptomic analyses identified the upregulated serine/cysteine protease inhibitor, SERPINB3 (squamous cell carcinoma antigen 1, SCCA1) in basal-like/squamous PDAC using discovery and validation approaches. Upregulation of SERPINB3 associated with decreased patient survival and a transcriptome profile indicative of the basal-like/squamous subtype. In human PDAC cell lines, SERPINB3 transgene expression enhanced their invasion capability. Moreover, upregulated expression of SERPINB3 in AsPC-1 cells resulted in enhanced lung metastasis in an orthotopic xenograft model. Molecular analysis of the primary tumor xenografts indicated activation of pathways related to metastasis, increased oxidative damage, and angiogenesis when SERPINB3 was upregulated. Furthermore, metabolomic analysis, using patient cohorts and PDAC cell lines showed a distinct metabolic pattern closely associated with both SERPINB3 and the basal-like/squamous subtype, which included upregulation of carnitine/acylcarnitine, amino acid, glutathione, and purine metabolic pathways, and glycolysis. Further RNA-seq and metabolomic analyses indicated that SERPINB3 may potentially induce the basal-like/squamous subtype and metabolic reprogramming through MYC activation. Taken together, our findings identified SERPINB3 as a candidate marker gene for the basal-like/squamous subtype, which may contribute to the disease aggressiveness in this subtype of PDAC.

**Abbreviations:** 8-Hydroxy-2’-deoxyguanosine (8-OHdG), Diaminobenzene (DAB), Gene Set Enrichment Analysis (GSEA), Human Metabolome Technologies, Inc. (HMT), Immunohistochemistry (IHC), Ingenuity pathway analysis (IPA), Pancreatic ductal adenocarcinoma (PDAC), Serine/Cysteine Proteinase Inhibitor Family B Member 3 (SERPINB3)

**Graphical abstract:** 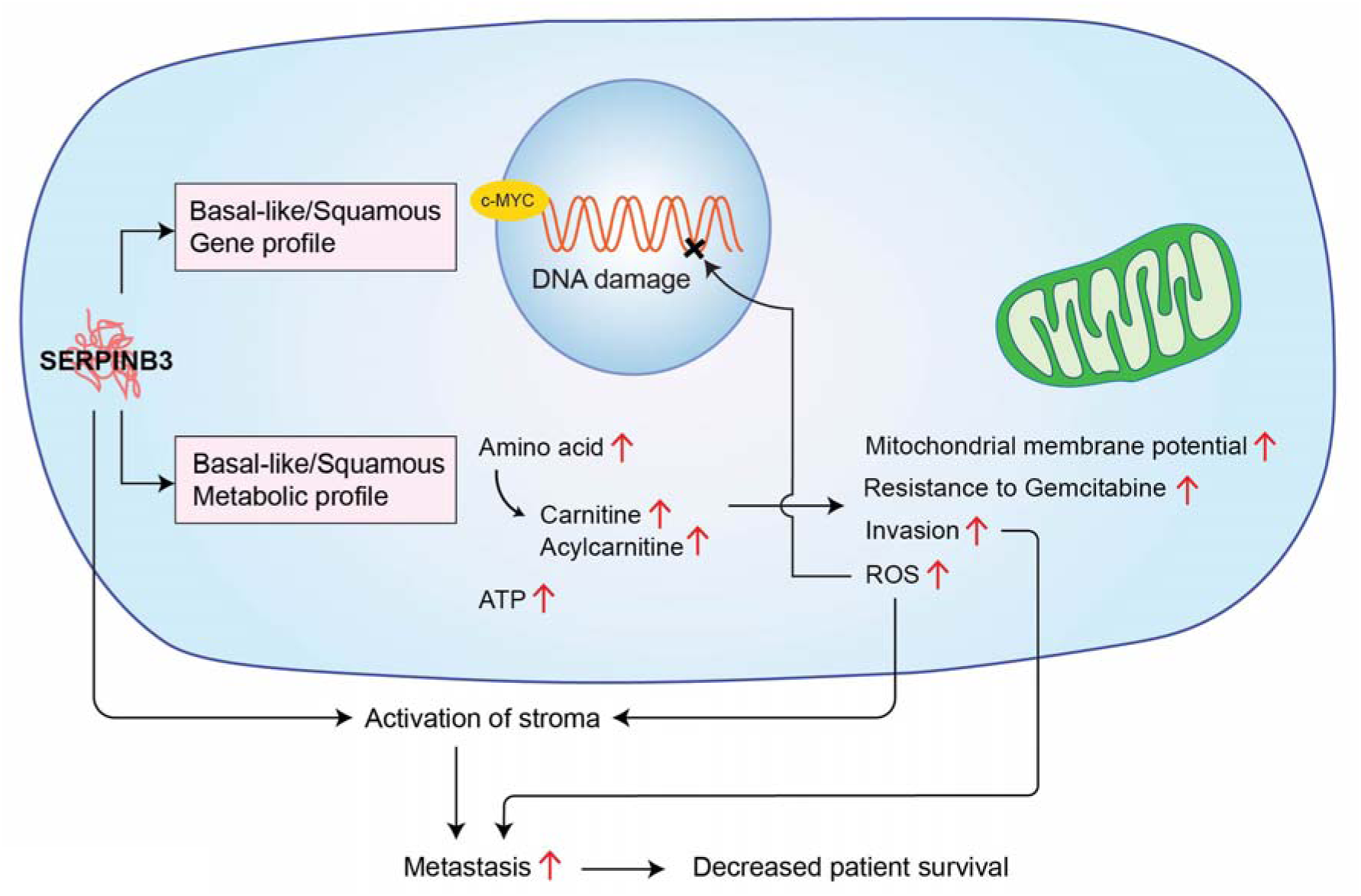

**Highlights:**

- SERPINB3 is upregulated in basal-like/squamous PDAC and associates with decreased patient survival
- SERPINB3 promotes differentiation into the basal-like/squamous subtype and enhances invasion and metastasis of PDAC
- SERPINB3 induces metabolic reprogramming and MYC activation and a metabolic signature indicative of basal-like/squamous PDAC

## Introduction

Pancreatic cancer is a lethal cancer with a 5 year survival of only 12% (1). Pancreatic ductal adenocarcinoma (PDAC) is the most common histologic form and comprises more than 90% of all malignancies in the pancreas (2).

Earlier studies identified several molecular subtypes of PDAC with differences in biological features and patient survival (3–5). *Moffitt et al.* proposed that PDAC can be separated into two subtypes; the basal-like and the classical subtype (4). Basal-like subtype contained characteristics similar to those of basal tumors in breast and bladder cancer. They also classified the stroma as ‘normal’ and ‘activated’. In contrast, *Bailey et al.* argued that PDAC can be classified into four subtypes: squamous, immunogenic, pancreatic progenitor, and aberrantly differentiated endocrine exocrine (3). The squamous subtype associated with the most aggressive disease and exhibited distinct features, including metabolic reprogramming, autophagy, inflammation, hypoxia, squamous differentiation, RNA processing, and activated MYC pathways (3). Recently, a comprehensive molecular subtype analysis using a transcriptome-based data approach classified PDAC into two subtypes; ‘classical/progenitor’ and ‘basal-like/squamous’ (6). The authors reported that the classical/progenitor subtype possesses an endodermal-pancreatic characteristic, while the basal-like/squamous subtype has lost this identity (6,7).

Metabolic reprogramming is reported as one of the hallmarks of cancer (8). Previous studies have described metabolic adaptations in PDAC as one of the key events in growth and progression (8,9). Cancer cells increase aerobic glycolysis, amino acid and lipid synthesis, and macromolecule synthesis to fulfill the high demand of energy and regulate oxidative stress (8–11). In terms of metabolic adaptations, PDAC may have a distinct metabolic profile (12–15). The classical/progenitor subtype may rely on lipogenic pathways for its metabolic needs whereas the basal-like/squamous subtype is more dependent on glycolysis (12–15). However, the association between metabolic profile and gene profile has not been well elucidated. These previous studies were based on cell lines or small number of PDAC patient samples. In this study, we have tested the hypothesis that an integrative transcriptomics and metabolomic approach may identify potential contributor of tumor progression and disease aggressiveness in the most aggressive basal-like/squamous subtype of PDAC, using large PDAC patient cohorts and human PDAC cell lines.

To test this hypothesis, we first analyzed the two large data sets (GSE36924 and GSE71729) (3,4) to filter the genes that were commonly associated with basal-like/squamous subtype in both data sets. We then validated these potential genes in our NCI-UMD-German cohort for their association with patient survival using Kaplan-Meier analysis. Furthermore, the validated genes were analyzed for their expression level in tumors and adjacent nontumor pancreas in our patient cohort. This integrative analysis resulted in the identification of serine/cysteine protease inhibitor family B member 3 (SERPINB3, also known as squamous cell carcinoma antigen 1, SCCA1) as a top candidate gene associated with poor patient survival in basal-like/squamous subtype, and therefore selected for further investigation of its functional role in disease progression. Furthermore, metabolomics and its integration with transcriptomic analyses were performed for mechanistic and functional role of SERPINB3 using our patient cohort, cell lines, and orthotopic mouse model. Our findings suggest that SERPINB3 contributes to the aggressiveness in basal-like/squamous subtypes through metabolic reprogramming and may be potentially targeted for designing improved treatment strategies.

## Materials and Methods

### Human samples

Pancreatic tumor tissues from resected PDAC patients were collected at the University of Maryland Medical System (UMMS) in Baltimore, MD, through an NCI-UMD resource contract and at the University Medical Center Göttingen, Germany. PDAC histopathology was determined by board-certified pathologists. The use of these clinical samples has been approved by the NCI-Office of the Human Subject Research Protection (OHSRP, Exempt#4678) at the NIH (Bethesda, MD).

### Public datasets

Transcriptome datasets derived from the “Bailey” cohort (GSE36924) (3) and “Moffitt” cohort (GSE71729) (4) were used for identifying genes of origin for the basal-like/squamous subtype, using the Partek Genomics Suite (Partek Inc., Chesterfield, MO) to generate gene signatures and to compare the basal-like/squamous subtype with the classical/progenitor subtype.

### Gene expression analysis

Microarray-based data from patient tumors: mRNA expression profiling was performed with Affymetrix GeneChip Human 1.0 ST arrays at the microarray core facility of the National Cancer Institute, Frederick, MD. The data was acquired and processed, as described earlier (16). The mRNA microarray expression data have been deposited to the National Center for Biotechnology Information’s (NCBI’s) Gene Expression Omnibus (GEO) database under accession number GSE183795.

RNA sequencing data from patient tumors: For bulk RNA sequencing on human PDAC patient samples, libraries were prepared by the Sequencing Facility at NCI-Leidos using the TruSeq Stranded mRNA Kit (Illumina, San Diego, CA) and sequenced paired-end on NovaSeq (Illumina) with 2 × 150 bp read lengths. About 51 to 122 million paired end reads in total were generated with a base call quality of Q30 and above. The sequence reads in fastq format were aligned to the human reference genome hg38 using STAR and RSEM to obtain gene expression as transcript per million and FPKM mapped reads. Differential expression was performed using DESeq2. The RNA-seq data from patient tumors were deposited in the NCBI’s GEO database under accession number GSE224564.

RNA sequencing data from PDAC cell lines: We performed quadruplicate RNA-sequencing on human PDAC cell lines (AsPC-1 ± SERPINB3; Panc 10.05 ± SERPINB3). Cells were cultured with regular media for 72 hours before RNA extraction. Libraries were prepared using the TruSeq Stranded mRNA Kit (Illumina) and sequenced paired-end on NextSeq (Illumina) with 2 × 150 bp read lengths. About 22 to 29 million paired end reads were generated with a base call quality of Q30 and above. Data analysis was processed, as described in human PDAC patient samples. The RNA-seq data for human PDAC cell lines were deposited in the NCBI’s GEO database under accession number GSE223908.

RNA sequencing data from tumor xenografts: RNA sequencing on the pancreatic xenografts in the orthotopic mouse model using PDAC cells (AsPC-1 ± SERPINB3) was also performed following isolation of RNA with the RNeasy Plus Mini Kit (QIAGEN, Venlo, Netherlands). Sequencing was performed on NovaSeq 6000 SP (Illumina) with 2 × 150 bp red lengths. About 133 to 164 million paired end reads in total were generated with a base call quality of Q30 and above. The sequence reads in fastq format were aligned to the human reference genome hg38 and the mouse reference genome mm10 using STAR and RSEM to obtain gene expression as transcript per million and FPKM mapped reads. Differential expression was performed using DESeq2. The RNA-seq data for the xenografts were deposited in the NCBI’s GEO database under accession number GSE223536.

### PDAC subtype assignment using the published “Moffitt” gene set

A list of signature genes was obtained as earlier described by *Moffitt et al.* (4). We obtained the expression pattern of the signature genes from RNA sequencing and assigned those to each tumor as a score and performed hierarchical clustering with Euclidean distance and the ward D2 clustering algorithm. The approach split the tumors into 3 main subtypes that we manually curated as classical/progenitor, basal-like/squamous, and unclassified.

### Quantitative real-time PCR for *SERPINB3*

The High-Capacity cDNA Reverse Transcription Kit (Applied Biosystems/Thermo Fisher Scientific, Waltham, MA) was used to create first-strand cDNA from total RNA. Quantitative RT-PCR (qRT-PCR) assays were performed using Taqman probes (Applied Biosystems/Thermo Fisher Scientific): *SERPINB3* (Hs00199468_m1) and *GAPDH* (Hs99999905_m1).

### Metabolic profiling and data analysis of tumor samples from PDAC patients

The metabolic profiling of our tumor samples from PDAC patients was conducted by Metabolon Inc. (Morrisville, NC) using a standard protocol, as previously described (17–20). For the current study, three existing metabolome profiling data sets (n = 33, n = 31, and n = 33) were combined and 235 common metabolites of known identity were analyzed. The merged data was normalized by assigning the median of each compound level to equal to one for each dataset termed “block normalization”. We imputed missing data with the minimum observed value for each metabolite in each dataset and the combined dataset was log2-transformed. Only patient samples with RNA sequencing and follow up data were included in this study (n = 88). The dataset served for further analysis (Supplementary Table 1).

### Metabolome analysis of PDAC cell lines

Absolute concentration of 116 metabolites was examined in various cell lines using the metabolome analysis package “Carcinoscope” provided by Human Metabolome Technologies, Inc. (HMT) (Boston, MA). For metabolite extraction, cell extracts were obtained using the standard manufacturer’s protocol as described previously (21). Briefly, 1 × 10^6^ cells were seeded on 100 mm dish and incubated with RPMI1640 with 10%FBS for 72 hours in incubator. After the removal of medium from a dish, 800 μl of methanol was added and incubated at room temperature for 30 sec. 550 μl of Internal Standard Solution were added and incubated at room temperature for 30 sec. One ml of the extracted solution was transferred to a 1.5 ml microtube and centrifuged at 4°C for 5 min (2300 × g). 350 μl of the supernatant was transferred into a centrifugal filter unit and centrifuged at 4°C for 5 hours (9100 × g). Filtered samples were stored at −80°C until shipping. Metabolome analysis was performed by HMT on SERPINB3-overexpressing PDAC cells (n = 4) and vector control PDAC cells (n = 4) using capillary electrophoresis mass spectrometry (CE-MS).

### Cell lines and culture condition

We purchased human PDAC cell lines from American Type Culture Collection (ATCC), Rockville, Maryland. Cell lines were authenticated by short tandem repeat (STR) analysis at ATCC, per request, and were mycoplasma-free. PDAC cells were grown in RPMI 1640 medium with GlutaMax^TM^, 10% FBS, and 1% penicillin–streptomycin in a humidified incubator containing 5% CO2 at 37 °C. All reagents for cell culture were purchased from Thermo Fisher Scientific (Waltham, MA).

### SERPINB3 overexpression after lentiviral infection

The SERPINB3 construct (EX-F0390-Lv103) and the corresponding empty vector control (EX-NEG-Lv103) were purchased from Genecopoeia (Rockville, MD). To obtain stable cell lines overexpressing SERPINB3, PDAC cells (AsPC-1, MIA PaCa-2, and Panc 10.05) were infected with lentiviral particles produced by transfecting 293T cells using the lentiviral expression vectors and the Lenti-Pac^TM^ HIV Expression Packaging system from Genecopoeia. The PDAC cells were selected with puromycin (4 μg/ml) to obtain stable clones.

### Cell proliferation assay

PDAC cells were seeded in a 96-well plate and the CCK-8/WST-8 assay was performed according to the manufacturer’s protocol (Dojindo Laboratories, Kumamoto, Japan), at 0, 24, 48, 72, and 96 hours after seeding. The absorbance was measured with SpectraMax^®^ ABS Plus microplate reader (Molecular Devices, San Jose, CA).

### Cell invasion assay

24-well Falcon^®^ Cell Culture Insert and Matrigel Matrix were purchased from Corning Inc. (Corning, NY). The membrane of the upper insert was coated applying 100 μl of Matrigel matrix coating solution (Matrigel matrix: coating buffer = 1:39) for 2 hours according to the manufacturer’s protocol. PDAC cells (AsPC-1 10×10^4^ cells, Panc 10.05; 10×10^4^ cells) in 500 μl serum-free RPMI 1640 with GlutaMax^TM^ were loaded into each upper insert, and 750 μl of RPMI 1640 with GlutaMax^TM^ and 10% FBS were added to the lower chamber. We counted PDAC cells that passed through the cell culture insert membrane after 48 hours of incubation in a humidified incubator containing 5% CO2 at 37 °C. For fixation and staining of the cells, 100% methanol (Thermo Fisher Scientific, Waltham, MA) and Crystal violet solution (MilliporeSigma, Burlington, MA) were used. To examine the effect of carnitine and hydroxyproline, either 1 mM of carnitine or 0.5 mM of hydroxyproline (final conc.) were added into the lower chamber. Carnitine and hydroxyproline were purchased from MilliporeSigma, Burlington, MA.

### Measurement of mitochondria membrane potential and reactive oxygen species

PDAC cells were seeded in a 60 mm dish at 10^6^ cells with regular media and cultured for 48 hours. To examine effects of L-carnitine, 1 mM carnitine (final conc.) was added to the culture media after PDAC cells were seeded. We used CellROX™ Deep Red Reagent (Thermo Fisher Scientific, C10422) for reactive oxygen species (ROS) measurement and MitoTracker™ Red FM (Thermo Fisher Scientific, M22425) for measurement of the mitochondrial membrane potential. Here, PDAC cells were stained with 5 μM of CellROX™ Deep Red or 100 nM of MitoTracker™ Red FM for 30 minutes at 37°C. After washing twice with PBS, we evaluated ROS or mitochondrial membrane potential using a SONY SA3800 flow cytometer (Sony Group Corporation, Tokyo, Japan). Data analysis was performed using FlowJo version 10.8.1.

### Orthotopic injection of AsPC-1 cells into the pancreas of mice for tumor growth

Animal experiments and maintenance conformed to the guidelines of the Animal Care and Use Committee at NCI and the American Association of Laboratory Animal Care. Each experiment was approved by ACUC at NCI, Frederick, MD. Briefly. 5×10^5^ of either SERPINB3-overexpressing or control AsPC-1 cells were orthotopically transplanted in the pancreas of NOD-SCID mice. The mice were euthanized at 5 weeks after transplantation. A complete necropsy was performed and the primary tumors were harvested and weighed. Paraffin-embedded sections of the pancreas, liver, and lungs were prepared and evaluated for any pathological changes including metastasis.

### Immunohistochemistry

Four μm thick paraffin-embedded sections were prepared for immunohistochemistry (IHC). Antigen retrieval was performed by heating the sections with microwave in Target Retrieval Solution, pH 9 (Agilent Technologies, Santa Clara, CA). The sections were incubated with primary antibodies overnight at 4°C. The following antibodies were used: SERPINB3 (Abcam, Cambridge, United Kingdom, ab154971; 1:500), CD31 (Abcam, ab28364; 1:100), 8-Hydroxy-2’-deoxyguanosine (8-OHdG) (Abcam, ab48508; 1:5000). Signals were amplified using the Dako EnVision+ System- HRP labelled polymer anti-mouse or rabbit antibody protocol. Color development was conducted with diaminobenzene (DAB, Agilent Technologies). For the human PDAC samples, the immunostaining of SERPINB3 was evaluated assigning the intensity and prevalence score as described previously (22,23). Briefly, the intensity was assigned a score of 0–3, representing negative, weak, moderate, or strong expression and distribution was given a score of 0–4, with <10%, 10–30%, >30–50%, >50–80% or >80% cells showing SERPINB3 expression. Then the overall IHC score was obtained by multiplying the intensity and distribution scores. The density of microvessels in the orthotopic tumor xenografts was assessed using CD31 expression in endothelial cells. Oxidative stress was assessed by the percentage of 8-OHdG-positive tumor cells in the xenografts.

### Immunoblotting

Cells were lysed with RIPA Lysis and Extraction Buffer (Thermo Fisher Scientific). Proteins were electrophoresed under reducing conditions on 4–15% polyacrylamide gels (Bio-Rad Laboratories, Inc., Hercules, CA) and then transferred onto a nitrocellulose membrane (Bio-Rad Laboratories, Inc.). This membrane was incubated for 60 min with SuperBlock™ Blocking Buffer (Thermo Fisher Scientific) at room temperature. Incubation with primary antibody was carried out overnight at 4°C. The following primary antibodies were used: SERPINB3 (Bio-Techne Corporation, Minneapolis, MN, MAB6528; 1:500) and β-Actin (MilliporeSigma, Burlington, MA, A5441, 1:2000). The membrane was incubated with secondary ECL anti-rabbit or anti-mouse IgG HRP-linked antibody (GE Healthcare, Pittsburgh, PA) for 1 hour at room temperature. Protein was visualized using a SuperSignal™ West Dura Extended Duration Substrate (Thermo Fisher Scientific).

### Statistical analysis

Data were analyzed using GraphPad Prism 9 (GraphPad Software, La Jolla, CA). Overall survival among PDAC patients was evaluated using the Kaplan–Meier method and log-rank test for significance testing. A *p* value < 0.05 with a two-sided test was considered significant.

## Results

### SERPINB3 is upregulated in basal-like/squamous subtype of PDAC patients and associates with poor survival

Our study was aimed to identify candidate driver genes involved in the development and progression of the basal-like/squamous subtype of PDAC. We initially analyzed transcriptome data from two PDAC cohorts (Bailey cohort and Moffitt cohort) (3,4), which identified 399 genes that were differentially expressed in basal-like/squamous subtype when compared with that of the classical/progenitor subtype (cutoffs: fold change > 1.5 or < −1.5, p < 0.05), with 131 of the genes being upregulated in the basal-like/squamous subtype (Figure 1A). The genes were further interrogated for their association with patient survival (hazard ratio > 1.5 using cox regression analysis, p < 0.05) in our NCI-UMD-German patient cohort, yielding a narrowed list of 47 genes (Supplementary Table 2). Among them, *SERPINB3* was ranked one of the top and robustly upregulated genes in the basal-like/progenitor subtype (Supplementary Table 2). SERPINB3 is also known as squamous cell carcinoma antigen (24–27), associates with hypoxia (28,29), and promotes MYC activation (30,31), cancer stemness (32,33), metabolic reprogramming (28), and inflammation (34–36). These functions overlap the characteristics of basal-like/squamous subtype (3), and we chose SERPINB3 for further investigation. *SERPINB3* transcripts were also upregulated in tumor tissues when compared with nontumor tissues, and the upregulation of SERPINB3 transcript levels associated with decreased patient survival in both the NCI-UMD-German cohort and a validation cohort (Moffitt cohort) (4) (Figure 1B-C). Notably, *SERPINB3* transcripts were undetectable in about one-third (31/92) of nontumor samples but measurable in 97% (123/127) of our NCI-UMD-German cohort patient tumors using quantitative RT-PCR assay. Furthermore, SERPINB3 protein was detected in both the nucleus and cytoplasm of tumor cells by immunohistochemistry (Figure 1D). Additionally, SERPINB3 protein upregulation in tumors also associated with a decreased survival in our NCI-UMD-German patient cohort (Figure 1D), consistent with the association of poor patient survival and enhanced SERPINB3 transcript levels. These findings are consistent with the hypothesis that SERPINB3 is associated with worst patient survival in basal-like/squamous subtype of PDAC and may be associated with aggressive disease progression.

**Figure 1:**
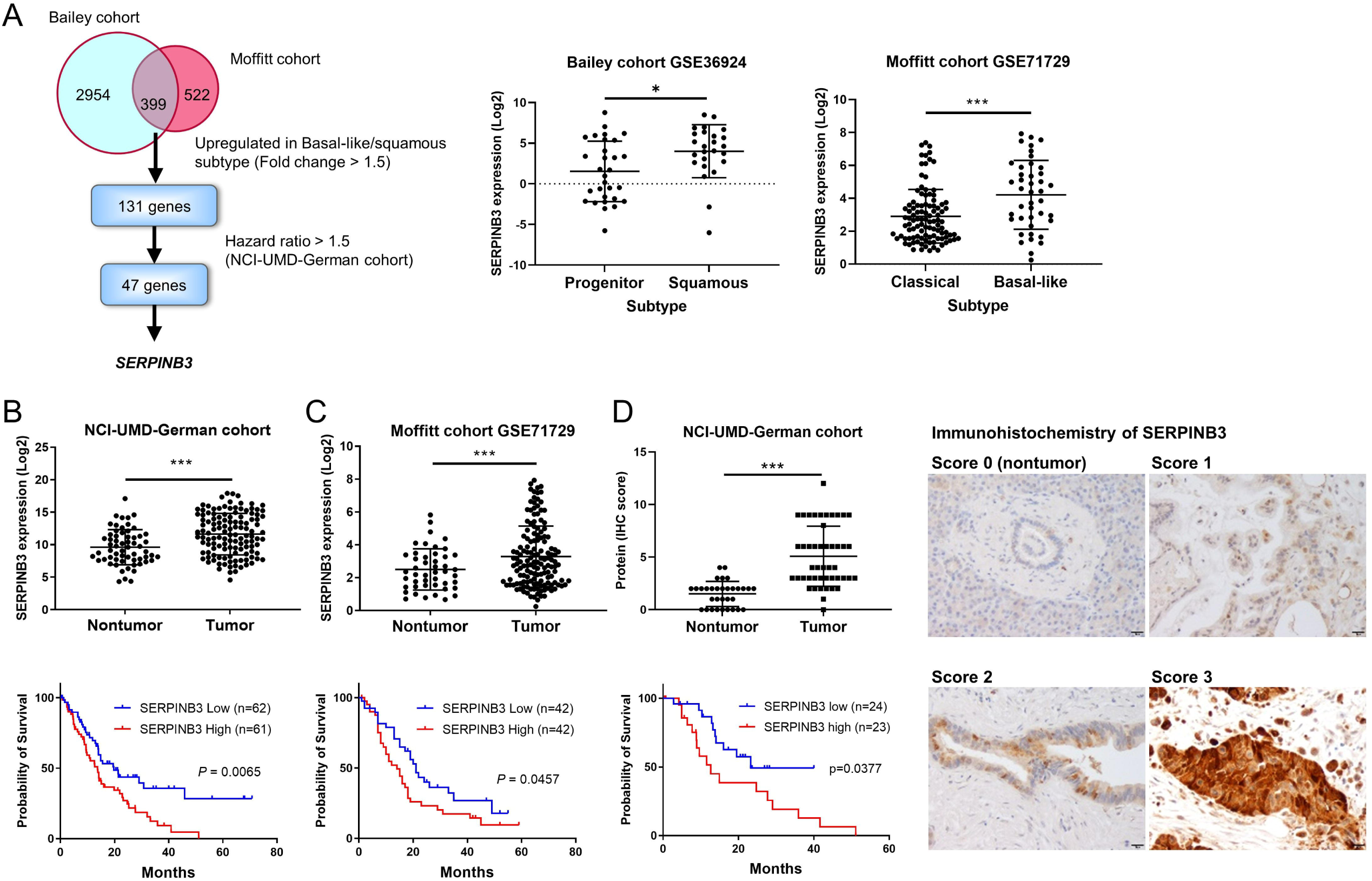
Transcriptome analysis identifies SERPINB3 as a marker upregulated in basal-like/squamous PDAC and associated with poor survival. (A) Strategy applied to find candidate driver genes of the basal-like/squamous subtype in PDAC. Two cohorts (Bailey and Moffitt cohorts) (3,4) were interrogated to identify candidate driver genes of the basal-like/squamous subtype. The genes were narrowed down to those associated with patient survival in the NCI-UMD-German cohort. (B-D) SERPINB3 transcript and protein expression in PDAC tumors versus adjacent non-cancerous tissues showing upregulation of SERPINB3 in tumors in both the NCI-UMD-German cohort and a validation cohort (Moffitt cohort; GSE71729) (only mRNA). The lower panel shows Kaplan-Meier plots and the association of increased SERPINB3 with decreased PDAC patient survival. For the survival analysis in the validation cohort, patients in the upper and lower tertiles of SERPINB3 expression were compared. The panel to the right shows immunohistochemical detection of SERPINB3 in representative nontumor (score 0) and tumor tissues (scores 1-3). SERPINB3 protein was detected in both the nucleus and cytoplasm, as shown by the brown DAB-based immunostaining in the tumor cells. The images show the staining strength (score 0, nonstained; score 1, weak; score 2, moderate; score 3, strong). Scale bar is 20 μm. More details can be found in Materials and Methods. Data represent mean ± SD. Significance testing with unpaired t-test. *p < 0.05, **p < 0.01, ***p < 0.005

### Upregulation of SERPINB3 promotes the characteristics of basal-like/squamous subtype of PDAC

Next, we investigated if SERPINB3 expression promotes basal-like/squamous subtype characteristics with enhanced disease aggressiveness. Earlier described gene signatures for molecular subtypes (4) separated 175 patient of our NCI-UMD-German cohort into unclassified, basal-like/squamous and classical/progenitor subtypes (Figure 2A). Tumors defined as basal-like/squamous subtype showed upregulation of *SERPINB3* mRNA expression and associated with worse patient survival (Figure 2A). In the pathway enrichment analysis using the Ingenuity Pathways Analysis (IPA) in the NCI-UMD-German cohort, both the SERPINB3-high and basal-like/squamous PDAC tumors showed similar pathway activation patterns, including increased cellular movement (Figure 2B and 2C) and MYC signaling, as indicated by an upstream regulator analysis (Figure 2B and 2C; Supplementary Table 3). Additional Gene Set Enrichment Analysis (GSEA) revealed that 374 MYC signature genes defined by their MYC binding sites (37,38) and a basal-like gene expression profile derived from human breast tumors (39) were significantly enriched in the transcriptome of SERPINB3-high and basal-like/squamous PDAC tumors (see enrichment plots in Figure 2B and 2C).

**Figure 2.**
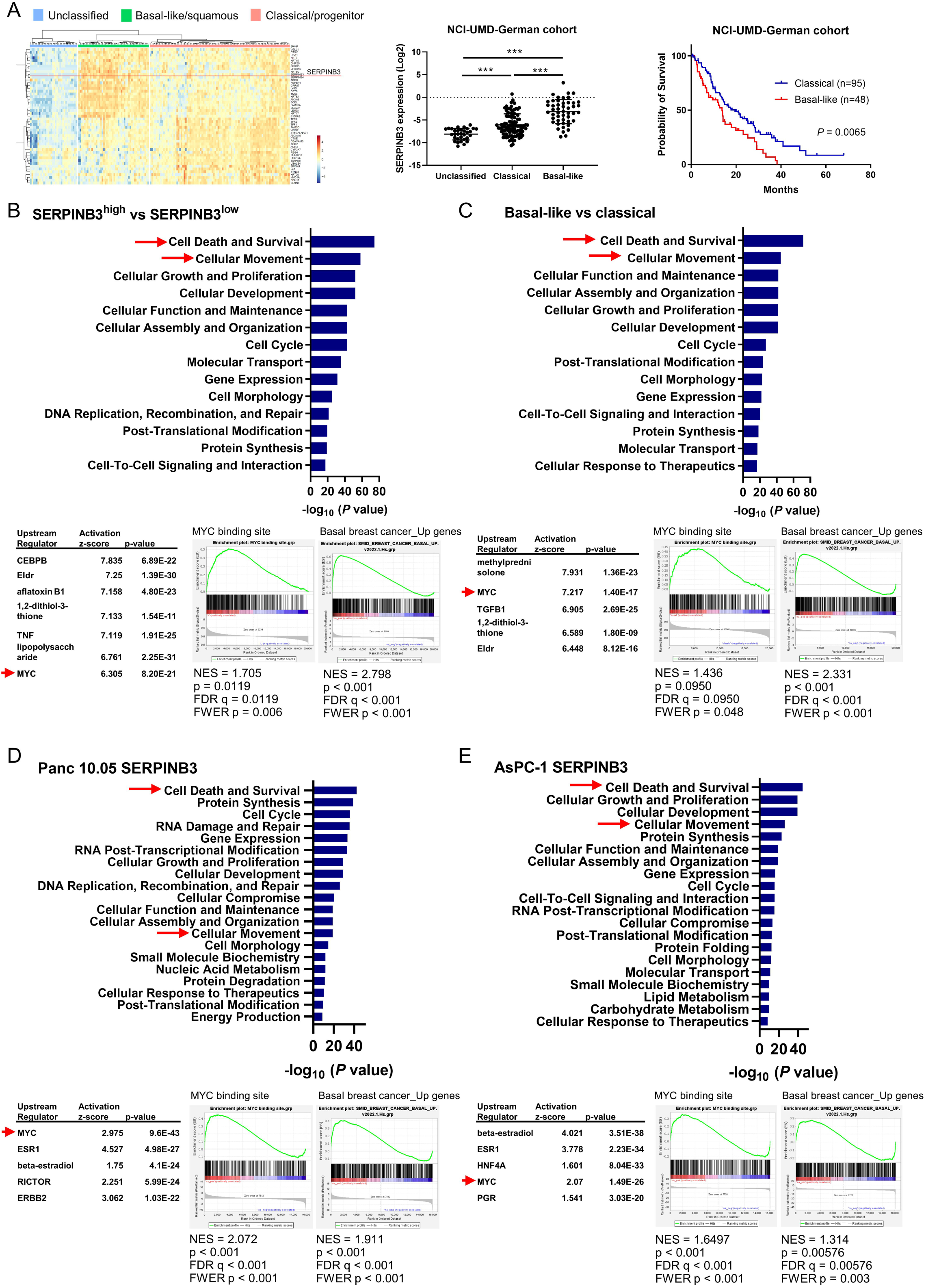
Upregulation of SERPINB3 promotes the characteristics of basal-like/squamous subtype of PDAC. (A) The transcriptome of PDAC defines molecular subtypes. Earlier described gene signatures for molecular subtypes (4) separate 175 patients of our NCI-UMD-German cohort into unclassified, basal-like/squamous, and classical/progenitor subtypes. Tumors defined as basal-like/squamous show upregulation of SERPINB3 mRNA expression (middle panel) and an association with inferior patient survival in the NCI-UMD-German cohort (right panel). (B-E) IPA (blue bars and tables) and GSEA highlight the activation of the similar pathways, including increased cellular movement, MYC activation, and induction of basal-like gene profile, in SERPINB3-high and basal-like/squamous PDAC tumors (B and C) and in human PDAC cell lines with SERPINB3 transgene overexpression (D and E). IPA; Ingenuity pathway analysis, GSEA; gene set enrichment analysis.

To follow up on our observations in patient tumors, we examined SERPINB3 expression levels in several human PDAC cell lines. Most of them expressed SERPINB3 (Supplementary Figure 1A). AsPC-1, MIA PaCa-2, and Panc 10.05 cells were then selected to establish cell lines with *SERPINB3* transgene overexpression. These cell lines are of the classical PDAC subtype (40), and can therefore be used to investigate the hypothesis that SERPINB3 overexpression potentially induce the transition of classical/progenitor subtype into the basal-like/squamous subtype. SERPINB3 overexpression was confirmed at the protein level in these cell lines (Supplementary Figure 1B). Pathway enrichment analyses using IPA and GSEA indicated a coherent activation of the same pathways across these human PDAC cell lines with *SERPINB3* transgene overexpression (Figure 2D and 2E; Supplementary Table 3), mirroring the findings from the SERPINB3-high and basal-like/squamous PDAC tumors. To elucidate SERPINB3 functions, additional *in vitro* experiments were performed using the SERPINB3-overexpressing PDAC cells. SERPINB3 overexpression did not have obvious effects on proliferation (data not shown). In contrast, SERPINB3 overexpression significantly enhanced the invasive ability of PDAC cell lines and reduced the sensitivity to standard of care drug gemcitabine (Figure 3A and 3B). These findings suggest that SERPINB3 contributes to maintain the characteristics of the basal-like/squamous subtype of PDAC.

**Figure 3.**
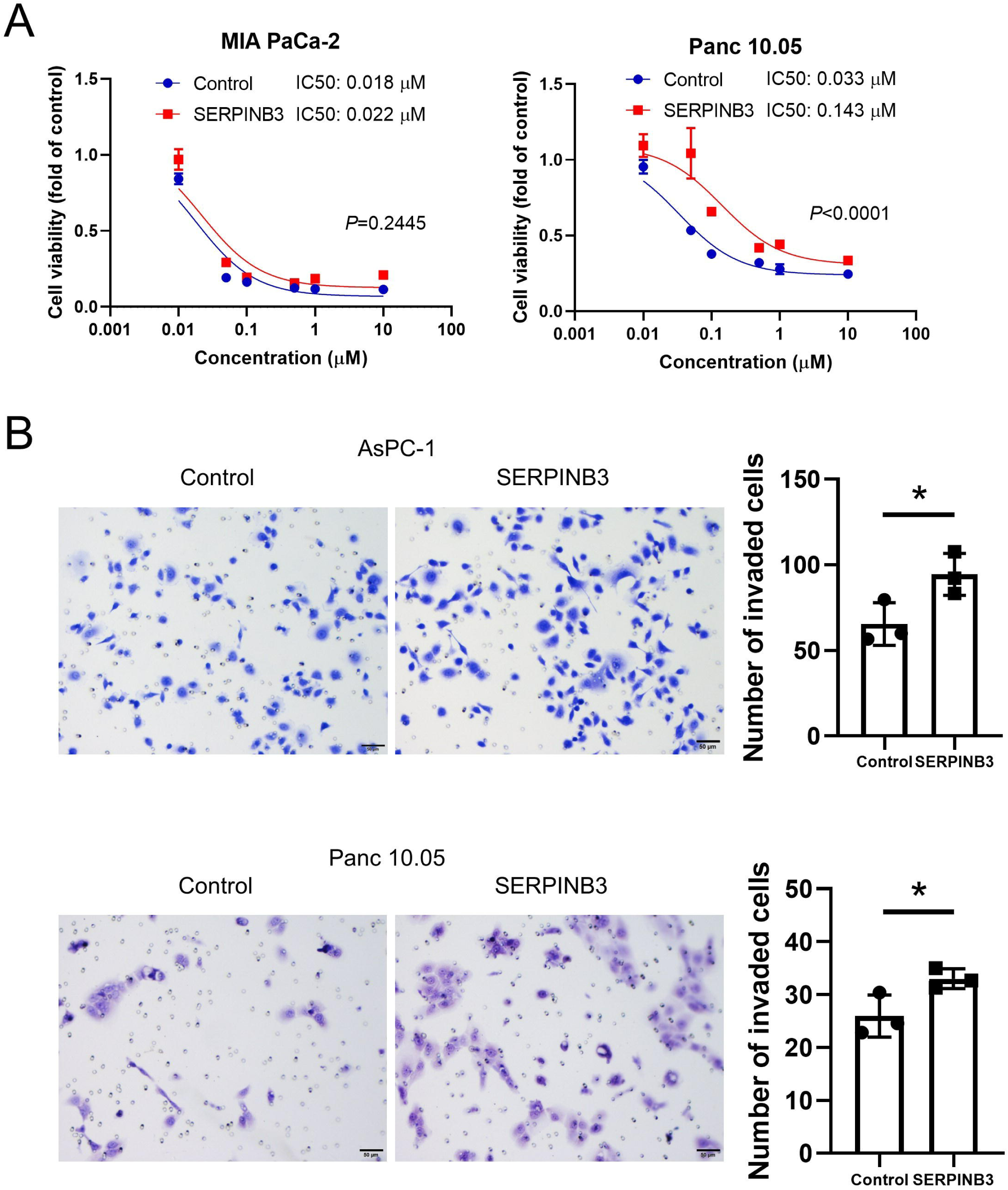
Upregulation of SERPINB3 promotes invasion of PDAC cells. SERPINB3 transgene expressing PDAC cells were examined to define the function of SERPINB3. (A) Dose-response curves showing viability of gemcitabine-treated PDAC cell lines (CCK-8/WST-8 assay). SERPINB3 reduces gemcitabine sensitivity in Panc 10.05 but not in MIA PaCa-2 cells. (B) SERPINB3-overexpressing PDAC cells have an increased ability to invade. Cells that passed through a cell culture insert membrane coated with Matrigel were fixed after 48 hours and cells were counted. Data represent mean ± SD of 3 replicates with t-test. *p < 0.05, **p < 0.01, ***p < 0.005.

### SERPINB3 promotes metastasis in an orthotopic mouse model of PDAC

To further investigate the mechanistic and functional role of SERPINB3 in tumor progression, we utilized an orthotopic mouse model. Implantation of SERPINB3 overexpressing AsPC-1 cells in the pancreas of mice resulted in significantly enhanced lung tumor metastasis (Figure 4A). However, Upregulation of SERPINB3 did not alter primary tumor growth of AsPC-1 cells, as indicated by the comparable tumor weights in the SERPINB3-expressing and control cells (Figure 4B). Furthermore, RNA-seq transcriptome and pathway analysis of the primary tumor xenografts revealed that upregulated SERPINB3 enhanced cellular movement and free radical scavenging (Figure 4C), which is consistent with our observations of patient tumors and cultured cells showing an increased invasion ability (Figure 3B) and oxidative stress (Supplementary Figure 2) when SERPINB3 is upregulated. IPA for the xenograft stroma exhibited the activation of pathways associated with HIF1α signaling, movement of tumor cells and endothelial cells, and angiogenesis (Figure 4D and 4E). Furthermore, immunostaining of endothelial marker CD31 and the reactive oxygen species, 8-OHdG, revealed that angiogenesis and oxidative stress are upregulated in the tumor xenografts of SERPINB3-overexpressing AsPC-1 in orthotopic model (Figure 4F and 4G). These findings revealed that SERPINB3 upregulation enhances tumor metastatic potential *in vivo*.

**Figure 4.**
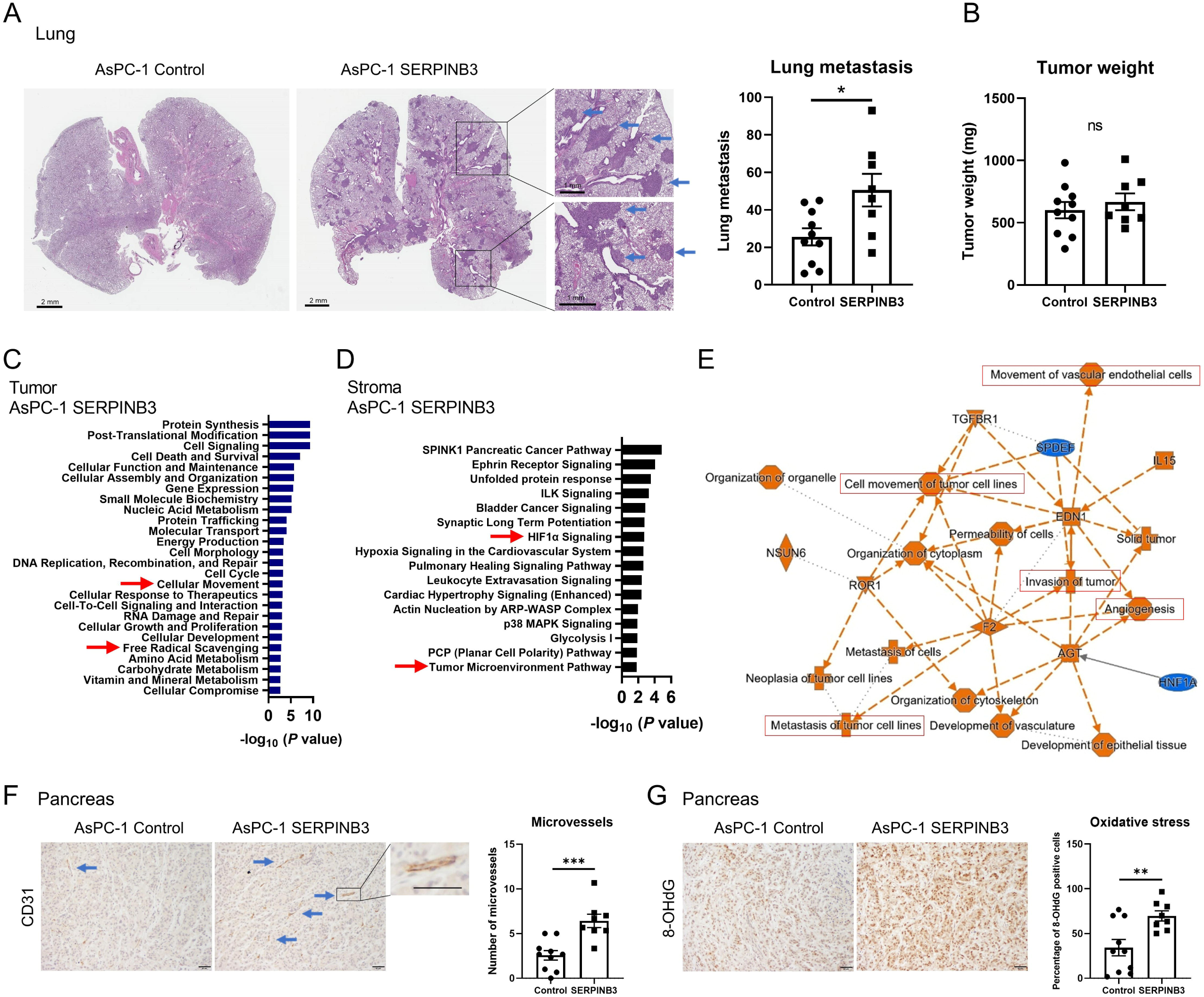
SERPINB3 promotes metastasis in an orthotopic mouse model of PDAC. Vector control and SERPINB3-overexpressing AsPC-1 human PDAC cells were transplanted into the pancreas of immune-deficient (NOD-SCID) mice. The tumor xenograft-growing mice were sacrificed at 5 weeks and both tumor burden and lung metastasis were assessed. (A) Increased number of lung metastasis in mice transplanted with AsPC-1 cells carrying the SERPINB3 transgene (shown in graph to the right). Basophilic clusters indicate the metastatic lesions in sections of the lung (see arrows). (B) Upregulation of SERPINB3 does not increase the weight of the primary tumor xenografts in the pancreas. (C) Pathway enrichment analysis using IPA for the primary tumor xenografts from SERPINB3 overexpressing AsPC-1 versus vector control cells. IPA indicates the activation of a set of pathways, including “cellular movement” and “free radical scavenging” in SERPINB3-overexpressing AsPC-1, consistent with increased invasion (Figure 3) and oxidative stress (Supplementary Figure 2) in these tumors. (D-E) Pathway enrichment analysis using IPA for the tumor stroma from xenografts of SERPINB3 overexpressing AsPC-1 versus vector control cells. The stroma in AsPC-1 SERPINB3 xenografts exhibits activation of pathways associated with “HIF1α” and “tumor microenvironment”, including “angiogenesis” and “metastasis/invasion/cell movement”. (F) The density of microvessels (angiogenesis) in the tumor xenografts examined by immunostaining for CD31. Microvessel density is increased in the stroma of SERPINB3-overexpressing AsPC-1 (arrows). Scale bar indicates 50 μm. (G) Immunostaining for 8-OHdG, a marker of oxidative stress, in the tumor xenografts. The percentage of the 8-OHdG positive cells as a measure for oxidative stress was assessed. The level of oxidative stress is increased in SERPINB3-overexpressing AsPC-1. Scale bar is 50 μm. Data represent mean ± SD. Significance testing with t-test. *p < 0.05, **p < 0.01, ***p < 0.005. IPA; Ingenuity pathway analysis.

### Upregulation of acylcarnitine/carnitine and amino acid metabolism in SERPINB3-high and basal-like/squamous PDAC

We next explored how SERPINB3 may influence the metabolism of PDAC by metabolic profiling of tumors from 88 patients in our NCI-UMD-German cohort (Supplementary Table 1). Thirty-two metabolites were significantly increased and 3 were decreased in SERPINB3-high tumors, when compared with SERPINB3-low tumors (Figure 5A and Supplementary Table 4). Furthermore, 15 metabolites were significantly increased and 2 were decreased in the basal-like/squamous subtype, when compared with the classical/progenitor subtype (Figure 5B, Supplementary Table 4). Among the 32 upregulated metabolites in *SERPINB3*-high tumors and the 15 upregulated metabolites in tumors of the basal-like subtype, 11 metabolites were common in both groups and included carnitine (a branched non-essential amino acid)/acylcarnitines (carnitine derivatives), amino acids, and carbohydrates (Figure 5C and 5D). Moreover, the upregulation of several acylcarnitines further associated with decreased patient survival (Figure 5C).

**Figure 5:**
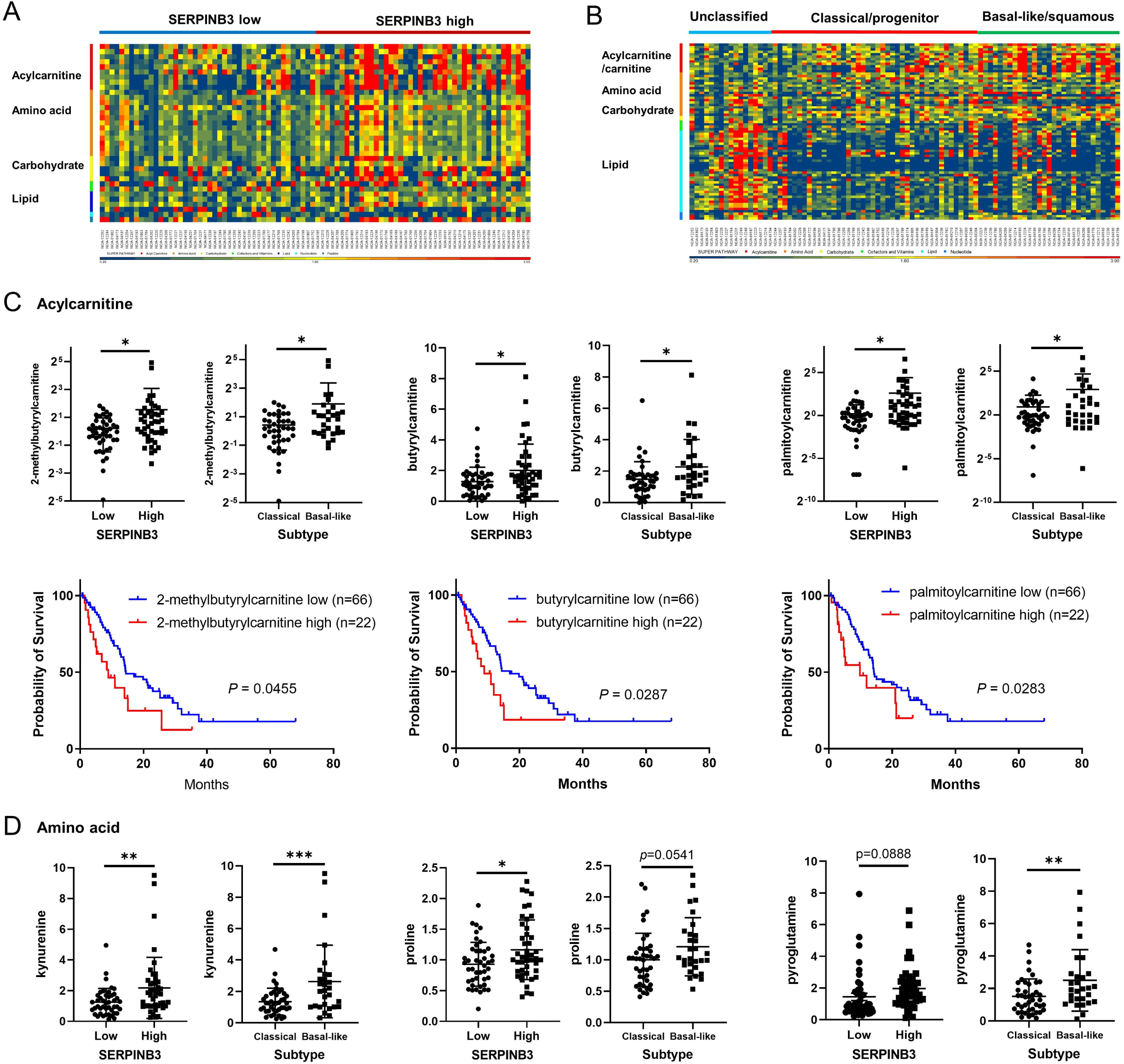
Upregulation of acylcarnitine/carnitine and amino acid metabolism in both SERPINB3-high and basal-like/squamous PDAC tumors. (A-B) Heatmaps show metabolite patterns in human PDAC (n = 88) with high and low SERPINB3 expression level (A) or by tumor subtype (B). Red color shows upregulation of metabolites, blue indicates low abundance. In A, 32 metabolites were significantly increased and 3 were decreased in SERPINB3-high tumors, when compared with SERPINB3-low tumors (p < 0.05). In B, 15 metabolites were significantly increased and 2 were decreased in the basal-like/squamous subtype, when compared with the classical/progenitor subtype (p < 0.05). Acylcarnitines/carnitine, amino acids, and carbohydrates are increased in SERPINB3-high PDAC tumors and also in basal-like/squamous PDAC tumors. (C-D) Abundance patterns for individual acylcarnitine (C) and amino acid (D) by tumor SERPINB3 status and subtype. The lower panel in C shows Kaplan-Meier plots depicting the association of each acylcarnitine with PDAC survival. A higher level of 2-methylbutylcarnitine, butyrylcarnitine and palmitoylcarnitine associated with poor patient survival. Data represent mean ± SD. Significance testing with t-test. *p < 0.05, **p < 0.01, ***p < 0.005.

To further investigate the role of these metabolites, we performed a global metabolome analysis of 116 cancer-related metabolites with absolute quantitation. In SERPINB3-overexpressing Panc 10.05 cells, 43 metabolites were increased when compared with control cells (Figure 6A and Supplementary Table 5). These metabolites represented the amino acid, glutathione, purine metabolism (e.g., ATP), and glycolysis/pentose phosphate pathway (PPP) (Figure 6B-C). Pathway enrichment analysis using MetaboAnalyst 5.0 (https://www.metaboanalyst.ca) with 43 input metabolites also indicated the upregulation of carnitine synthesis in SERPINB3-overexpressing Panc 10.05 cells (Figure 6C and Supplementary Table 5). In agreement with the transcriptome analysis, an upstream regulator analysis using the 43 metabolites as input for IPA predicted MYC signaling to be involved as a key regulator of metabolism in these cells as well (Figure 6D and Supplementary Table 6). MYC signaling was the only downstream regulator of SERPINB3 that was predicted by both our metabolome and transcriptome analysis (Figure 2). Furthermore, these findings were largely replicated in SERPINB3-overexpressing AsPC-1 and MIAPaCa-2 cells (Supplementary Figure 3, Supplementary Table 5, and Supplementary Table 6). These data showed that both basal-like/squamous subtype PDAC and SERPINB3-high cells have similar metabolic adaptation profile that may potentially contribute to the disease progression.

**Figure 6:**
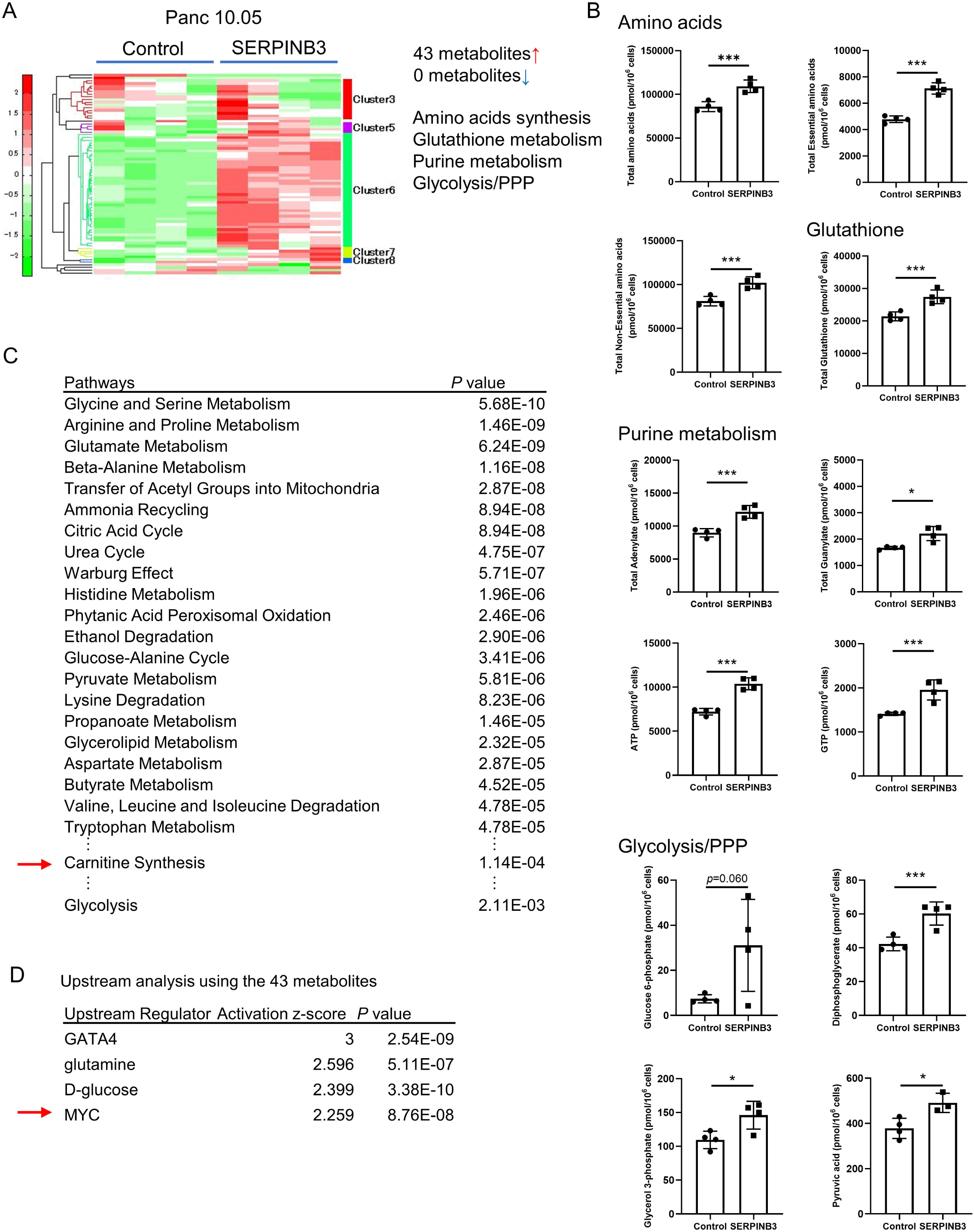
Upregulation of carnitine and amino acid metabolism in SERPINB3-overexpressing PDAC cells. (A-B) Metabolome analysis covering 116 cancer-related metabolites in SERPINB3-overexpressing Panc 10.05 cells. In SERPINB3-overexpressing Panc 10.05 cells, 43 metabolites were significantly elevated when compared with control cells (p < 0.05). Those represent metabolites for amino acids synthesis, glutathione metabolism, purine metabolism (e.g., ATP), and glycolysis/pentose phosphate pathway (PPP). (C) Pathway enrichment scores applying MetaboAnalyst 5.0 with 43 input metabolites. These metabolites were significantly increased in SERPINB3-overexpressing Panc 10.05 cells. (D) Prediction of upstream regulators for the metabolism in SERPINB3-overexpressing Panc 10.05 cells. The upstream analysis by IPA used the 43 input metabolites and predicts MYC signaling to be involved as a regulator of metabolism in these cells. Data represent mean ± SD of 4 replicates with t-test. *p < 0.05, **p < 0.01, ***p < 0.005. IPA; Ingenuity pathway analysis.

### SERPINB3-associated metabolism enhances disease progression in PDAC

Based on the metabolomic profile from the patient tumors and PDAC cells, we focused on carnitine/acylcarnitines and amino acids for further functional investigations. We hypothesized that the sum of carnitine and acylcarnitines in the dataset (total carnitine) may reflect the function of carnitine/acylcarnitine more comprehensively than each of the carnitine/acylcarnitine. With this approach, we found that total carnitine was significantly increased in *SERPINB3*-high tumors and the basal-like/squamous subtype (Figure 7A and 7B). Moreover, a higher total carnitine associated with decreased patient survival (Figure 7C). Exposing pancreatic cancer cells to L-carnitine showed that it did not have any effect on proliferation, however, L-carnitine resulted in enhanced invasiveness (Figure 7D and 7E), further supporting our observation that SERPINB3 may enhance the metastatic process more so than the growth of primary PDAC tumors. L-carnitine treatment of PDAC cells resulted in increased mitochondrial membrane potential (Figure 7F), which is consistent with previous observations (41), and a potential mechanism for an increase in reactive oxygen species (ROS) in SERPINB3 expressing PDAC cells. Accordingly, SERPINB3-overexpressing PDAC cells also showed a higher mitochondrial membrane potential (Supplementary Figure 4) and produced increased amount of ROS, as shown in Supplementary Figure 2. Moreover, proline/hydroxyproline was distinctively increased in *SERPINB3*-high PDAC tumors, the basal-like/squamous subtype, and SERPINB3-overexpressing PDAC cells (Figure 5D and Supplementary Figure 5A), therefore, we examined if hydroxyproline may alter the phenotype of PDAC cells. Hydroxyproline did not show any effect on proliferation of PDAC cell lines, however, hydroxyproline increased invasiveness (Supplementary Figure 5B and 5C). These findings indicate that SERPINB3-induced metabolic alteration may have a potential role in tumor progression and disease aggressiveness in basal-like/squamous subtype.

**Figure 7.**
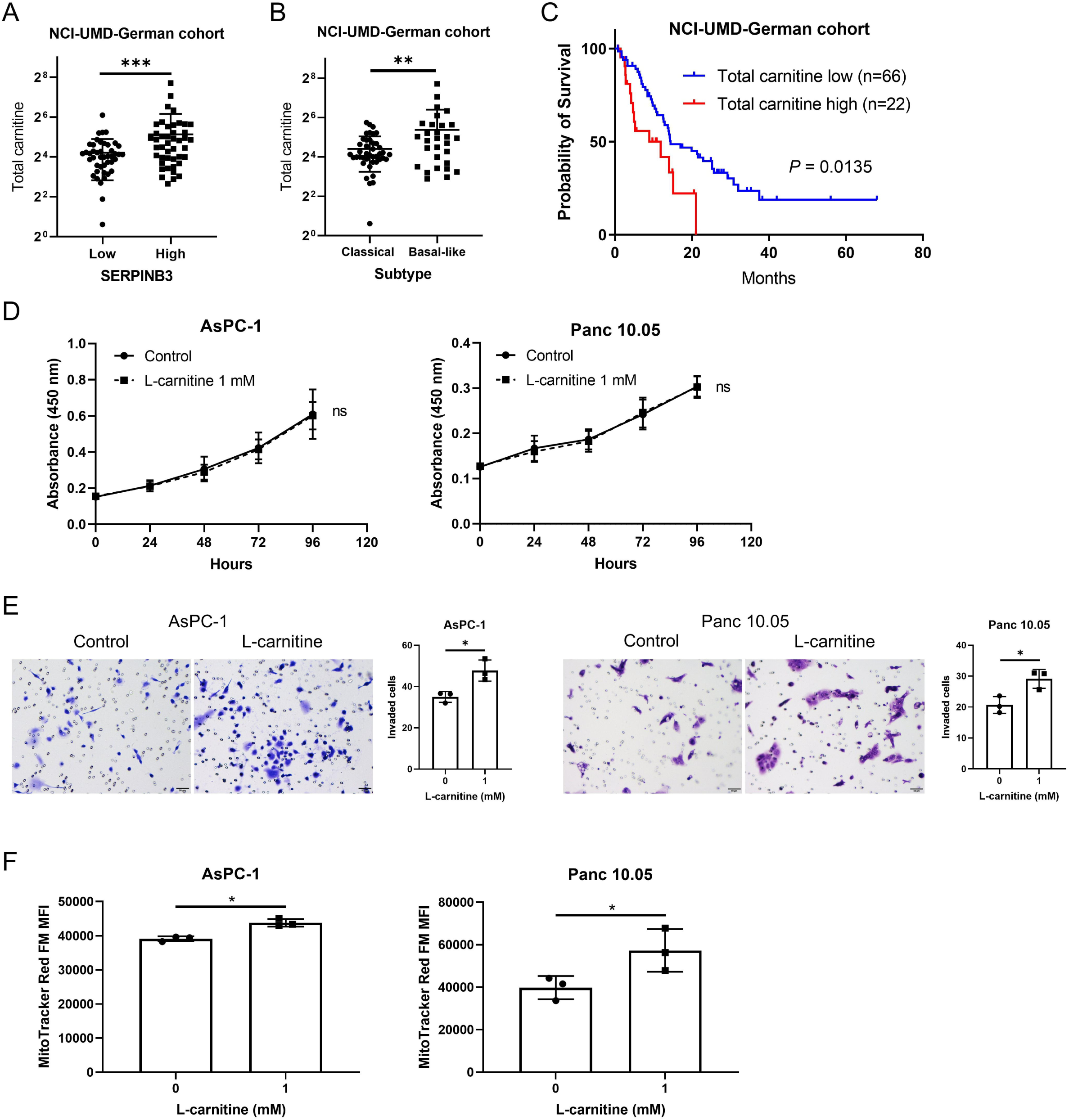
Carnitine enhances the disease progression in PDAC. (A-B) Total carnitine is elevated in both SERPINB3-high and basal-like/squamous PDAC in the NCI-UMD-German cohort. (C) Kaplan-Meier plot showing that high total carnitine associates with poor survival of PDAC patients (the upper 25% quartile vs the lower 75%; Total carnitine low < 30.8, high ≥ 30.8). (D) L-carnitine has no effect on proliferation of PDAC cell lines (CCK-8/WST-8 assay). (E) L-carnitine increases invasiveness of PDAC cell lines. Cells that passed through a cell culture insert membrane coated with Matrigel were fixed after 48 hours of incubation. (F) Increased mitochondrial membrane potential in PDAC cell lines cultured with 1 mM of L-carnitine for 48 hours. Mitochondrial membrane potential was quantified in cells using MitoTracker Red FM and flow cytometry. Data represent mean ± SD. Student t-test was used to determine the significance in difference. *p < 0.05, **p < 0.01, ***p < 0.005. Total carnitine is the sum of all acylcarnitines/carnitine in the metabolome dataset: 2-methylbutyrylcarnitine (C5), acetylcarnitine, butyrylcarnitine, carnitine, decanoylcarnitine, deoxycarnitine, hexanoylcarnitine, isobutyrylcarnitine, isovalerylcarnitine, octanoylcarnitine, oleoylcarnitine, palmitoylcarnitine, propionylcarnitine, stearoylcarnitine, succinylcarnitine, valerylcarnitine

## Discussion

Recently, a highly comprehensive analysis proposed that PDAC should be classified into just two subtypes; ‘classical/progenitor’ and ‘basal-like/squamous’ (6). However, none of previous studies identified key genes that promote different subtype characteristics including metabolic reprogramming, tumor progression, and disease aggressiveness. In the present study, we report that SERPINB3 is upregulated in the basal-like/squamous PDAC subtype and associated with worst patient survival. Furthermore, in PDAC preclinical model, SERPINB3 increased invasion and metastasis. Molecular analysis of primary tumor xenografts showed upregulation of pathways related to angiogenesis, oxidative stress, and metastasis. Mechanistic and functional analyses, primarily focused on metabolic alterations in patient tumors and cell line, showed an upregulation of glycolysis, carnitine, amino acid, glutathione, and purine metabolic pathways. Furthermore, our findings provided evidence that these SERPINB3-induced metabolic reprogramming is regulated by MYC activation. Taken together, these findings suggest that SERPINB3 contributes to the aggressive disease progression in the basal-like/squamous PDAC subtype.

Serine protease inhibitors (serpins) are one of the largest superfamilies of protease inhibitors with roles in regulating important pathways, such as coagulation and inflammation. Although most of the serpins are serine protease inhibitors, some also inhibit caspases (42) and papain-like cysteine proteases, including cathepsin L, S, and K, and papain (43,44). Serine/cysteine protease inhibitor family B member 3 (SERPINB3, also known as squamous cell carcinoma antigen 1, SCCA1) was initially described as a tumor-specific antigen in squamous cell carcinoma of the uterine cervix (24). SERPINB3 was later found to be upregulated in squamous cell carcinoma of the various organs (25–27) and other cancer, including hepatocellular carcinoma (34), cholangiocarcinoma (32), PDAC (36), glioblastoma (33), and breast cancer (35,45). In cancer, SERPINB3 has various functions including inhibition of radiation/anti-cancer drug-induced apoptosis, protein degradation, and intratumor infiltration of natural killer cells (27,34,46), and to promote metabolic reprogramming, angiogenesis, epithelial-mesenchymal transition, and secretion of inflammatory mediators (28,32–36). SERPINB3 also upregulates MYC (30,31), which leads to the maintenance of cancer stemness (32,33). Consistent with these findings, we found that SERPINB3 alters metabolism and induces a basal-like/squamous phenotype by the mechanisms that include MYC signaling pathway. In the orthotopic mouse model of PDAC, SEPRINB3 led to the activation of stroma with upregulation of angiogenesis and tumor microenvironmental pathways. Thus, SERPINB3 may play a role not only in the development and maintenance of the basal-like/squamous subtype of PDAC but also in inducing “activated” stroma, as described previously (4).

Cancer cells increase aerobic glycolysis, amino acid and lipid synthesis, and macromolecule synthesis using the upregulation of the pentose phosphate pathway (10,11). Genetic alterations are one of the drivers of such metabolic adaptations and may include amplification of the *MYC* locus. These adaptations result in appropriate maintenance of redox homeostasis, ATP generation, and promotion of their survival. MYC plays a central role of the activation of glycolysis and amino acid metabolism (11). MYC activates the transcription of glycolytic genes. MYC also increases the availability of essential amino acids through induction of amino acid transporters, such as SLC1A5, SLC7A5, and SLC43A1. Upregulation of tryptophan uptake leads to the kynurenine pathway. For the metabolisms of non-essential amino acids, including glutamine, proline, serine, and glycine, MYC promotes the biosynthesis/transport by upregulating the synthetic enzymes/transporters. Our findings showed that increased SERPINB3 expression results in a significant upregulation of glycolysis and most of the essential and non-essential amino acids metabolism (Supplementary Table 5) in cell lines. Consistently, similar association of metabolic adaptation was found under SERPINB3-high level in patient tumors of the basal-like/squamous subtype. Furthermore, upstream regulator analysis predicted the activation of MYC as a potential mechanism leading to these metabolic alterations. MYC may upregulate carnitine metabolism (11,47–49). Notably, MYC was found to upregulate carnitine uptake in triple-negative breast cancer (47), a representative type of basal-like breast cancer. In agreement with this finding and the candidate role of MYC as a regulator, downstream of SERPINB3, we observed the coherent up-regulation of carnitine/acylcarnitine metabolism in the basal-like/squamous subtype of PDAC or when SERPINB3 was upregulated. Thus, upregulation of the carnitine/acylcarnitine metabolism pathway might be one of the features of basal-like subtype tumors across the cancer spectrum.

Carnitine, a branched non-essential amino acid, is derived from the two amino acids, lysine and methionine (49–51). The function of L-carnitine is importing fatty acids into the mitochondria for use in β-oxidation by forming acylcarnitine derivatives from L-carnitine and acyl CoA, derived from fatty acids. As such, the endogenous carnitine pool is comprised of L-carnitine and various acylcarnitines. Carnitine also has a role in ejecting excess carbons (acetyl CoA) in the form of acylcarnitines from mitochondria, leading to glucose utilization and metabolic flexibility (50,52). Acylcarnitine is also involved in the metabolism of branched-chain amino acids (51). Several studies showed that carnitine increases mitochondrial membrane potential (41) and resistance against oxidative stress (53,54). The upregulation of carnitine/acylcarnitine may lead to a high energy supply using β-oxidation, increasing the mitochondrial membrane potential and ATP synthesis, as observed in the SERPINB3-overexpressing PDAC cell lines. Although the upregulation of carnitine/acylcarnitine metabolism in cancer has previously been documented, together with a role of carnitine/acylcarnitine metabolism in cancer progression (49-51,54,55), in our knowledge our study is the first to show its potential role in PDAC.

In summary, SERPINB3 contributes to the development and progression of the basal-like/squamous subtype of PDAC which is mediated by metabolic reprogramming through SERPINB3-MYC axis, which may be potentially useful in designing novel treatment strategies.

## Acknowledgments

Authors would like to thank all the staff of University of Maryland School of Medicine and NCI-University of Maryland study coordinators involved in the procurement of clinical biospecimens from pancreatic cancer patients. We would also like to thank the staff of the Department of General, Visceral and Pediatric Surgery, University Medical Center Göttingen, Göttingen, Germany for their help with clinical samples. We would also like to thank all the staff of the NCI-CCR Sequencing Facility Frederick, the NCI Laboratory Animal Sciences Program Frederick, and the CCR/LGI Flow Cytometry Core.

## Availability of data and materials

Data and materials are available upon request. The RNA-seq data are deposited in the NCBI’s Gene Expression Omnibus (GEO) database under accession number (GSE223909). To protect the confidentiality of our data and results, the repository will remain private while the manuscript is under review.

GSE223909: This SuperSeries is composed of the following SubSeries:

<u>GSE223536</u> SERPINB3 induces the aggressive basal-like/squamous subtype and promotes invasion and metastasis in pancreatic cancer I

<u>GSE223908</u> SERPINB3 induces the aggressive basal-like/squamous subtype and promotes invasion and metastasis in pancreatic cancer II

<u>GSE224564</u> SERPINB3 induces the aggressive basal-like/squamous subtype and promotes invasion and metastasis in pancreatic cancer III

To review GEO accession GSE223909:

Go to https://www.ncbi.nlm.nih.gov/geo/query/acc.cgi?acc=GSE223909 Enter token kjodywckzryrdqv into the box

All the metabolic data supporting the results in this article are included as the supplementary tables.

## Authors’ Contributions

**Conception and design:** Yuuki Ohara, S. Perwez Hussain

**Development of methodology:** Yuuki Ohara, Wei Tang, Shouhui Yang, Stefan Ambs, S. Perwez Hussain

**Acquisition of data (provided animals, acquired and managed patients, provided facilities, etc.):** Yuuki Ohara, Azadeh Azizian, Jochen Gaedcke, B. Michael Ghadimi, Nader Hanna

**Analysis and interpretation of data (e.g., statistical analysis, biostatistics, computational analysis):** Yuuki Ohara, Wei Tang, Huaitian Liu, Shouhui Yang

**Writing, review, and/or revision of the manuscript:** Yuuki Ohara, Wei Tang, Huaitian Liu, Shouhui Yang, Stefan Ambs, S. Perwez Hussain

**Administrative, technical, or material support (i.e., reporting or organizing data, constructing databases):** Yuuki Ohara, Wei Tang, Huaitian Liu, Shouhui Yang, Tiffany H. Dorsey, Helen Cawley, Paloma Moreno, Azadeh Azizian, Jochen Gaedcke, B. Michael Ghadimi, Nader Hanna

**Study supervision:** S. Perwez Hussain

## Supplementary figure legends

**Supplementary Figure 1.**
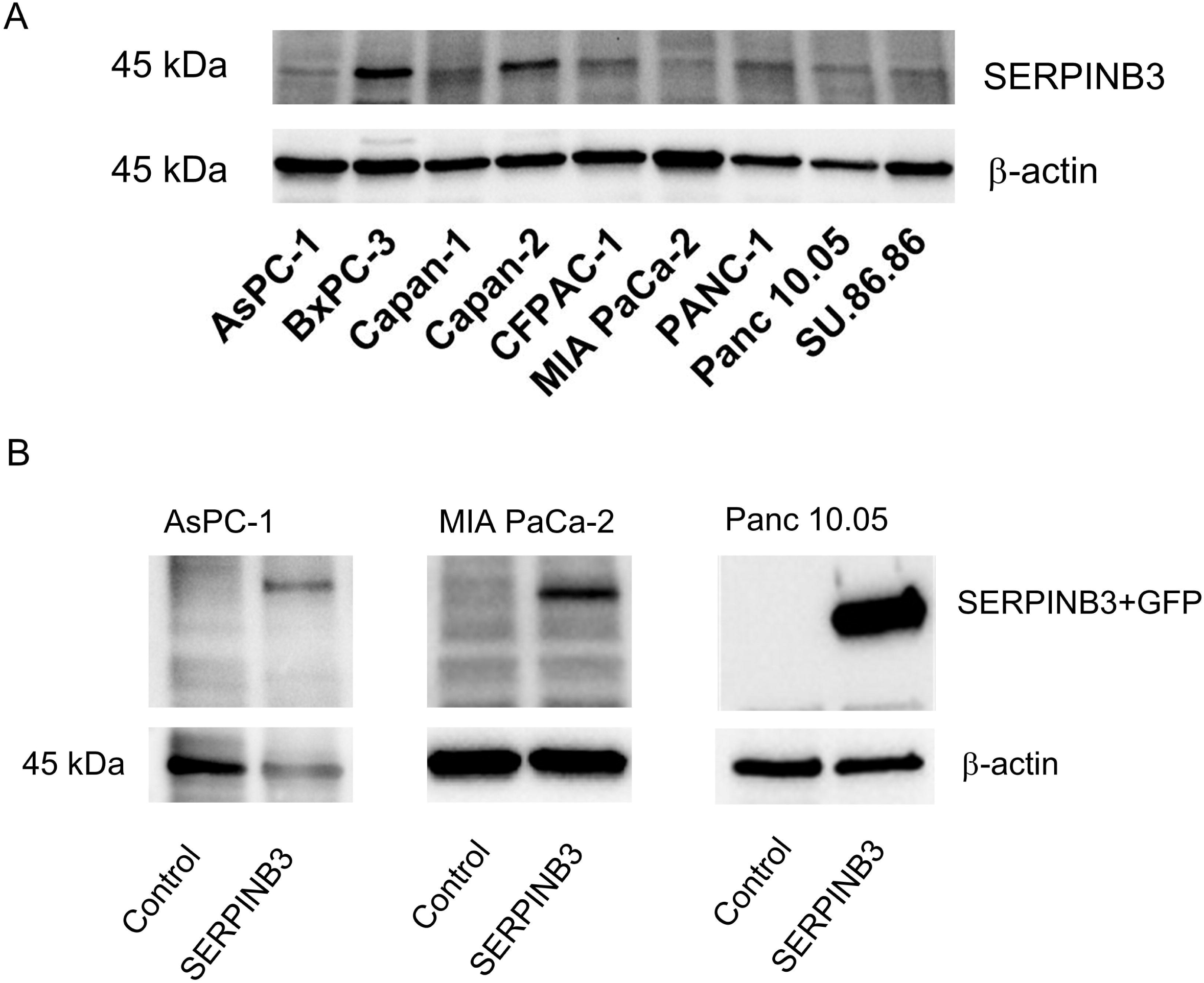
SERPINB3 expression in human PDAC cell lines. (A) Endogenous SERPINB3 protein levels in human PDAC cell lines. AsPC-1, MIA PaCa-2, and Panc 10.05 cells were selected to establish cell lines with SERPINB3 transgene expression. These cell lines have the classical PDAC subtype, as reported previously (40), and can be used to investigate the transition into the basal-like/squamous subtype after SERPINB3 overexpression. (B) Confirmation of SERPINB3 transgene overexpression at the protein level in AsPC-1, MIA PaCa-2, and Panc 10.05 cells.

**Supplementary Figure 2.**
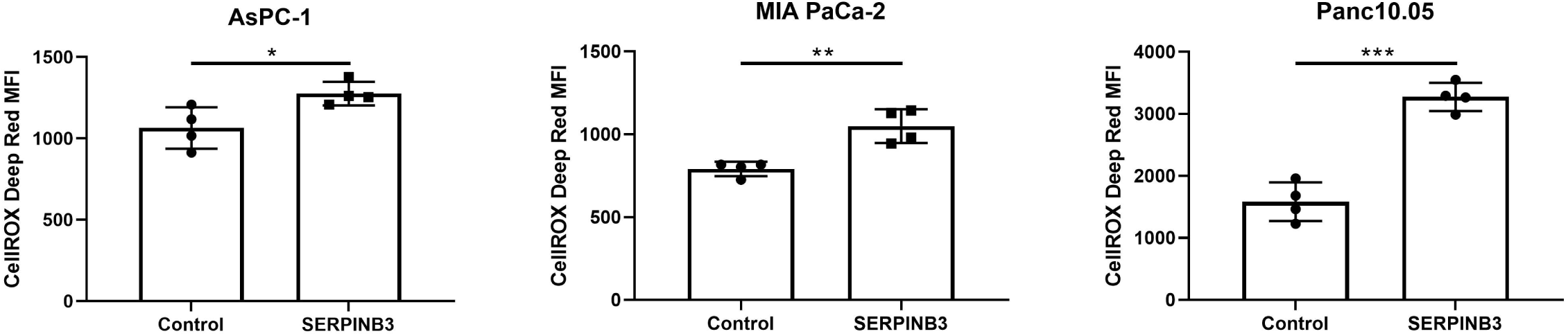
SERPINB3-overexpressing PDAC cells increase levels of reactive oxygen species *in vitro*. Levels of reactive oxygen species in the SERPINB3-overexpressing PDAC cells and the control PDAC cells, showing that SERPINB3 promotes oxidative stress levels. Data represent mean ± SD with t-test. *p < 0.05, **p < 0.01, ***p < 0.005. Reactive oxygen species level was measured with the CellRox Deep Red fluorogenic reagent.

**Supplementary Figure 3.**
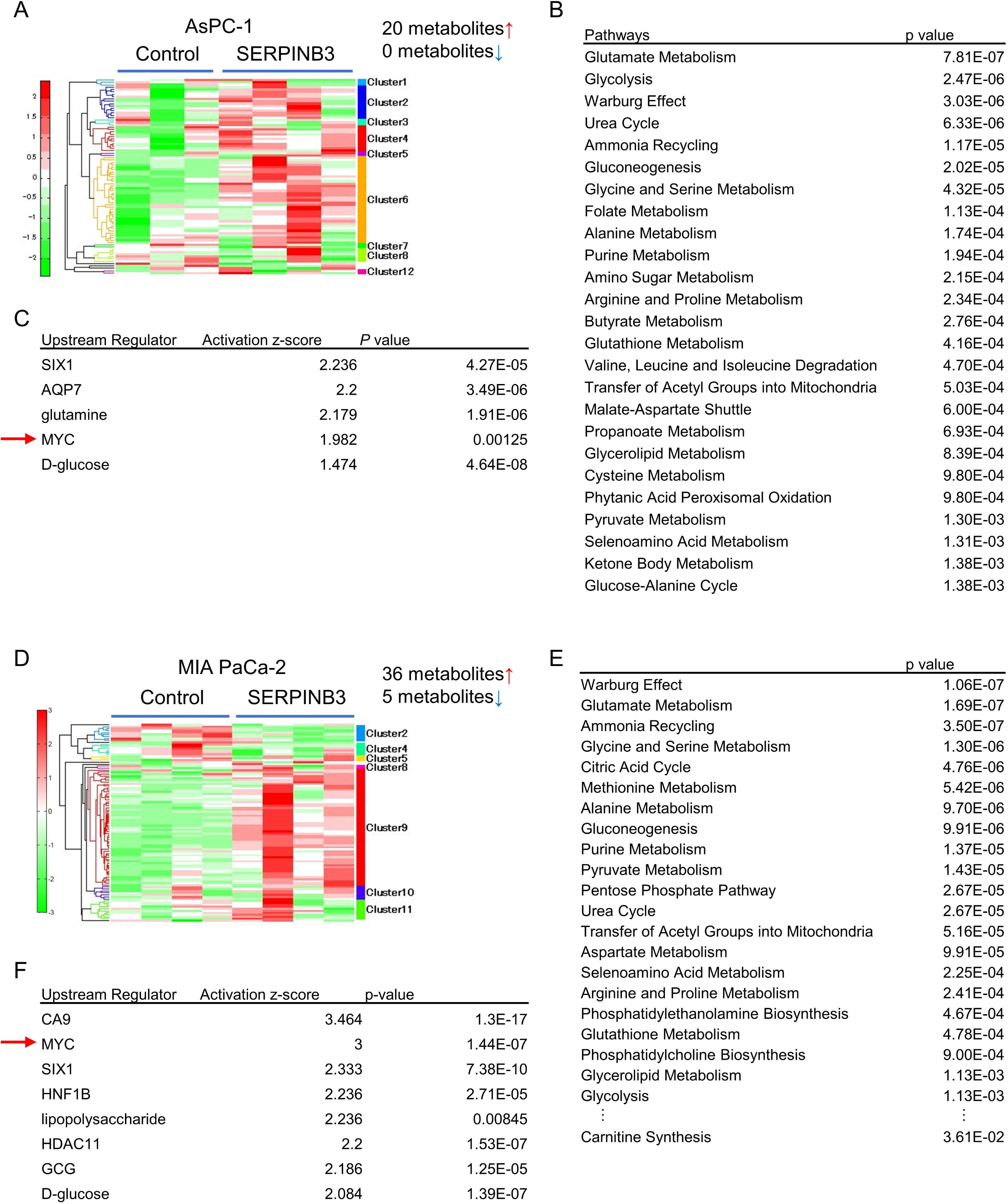
Upregulation of glycolysis and the Warburg effect and amino acid metabolism in SERPINB3-overexpressing PDAC cells. (A) Heatmap summarizing a metabolome analysis of 116 cancer-related metabolites in SERPINB3-overexpressing AsPC-1 cells, comparing their levels with vector control cells. 20 metabolites were significantly elevated in presence of high SERPINB3 (p < 0.05). Red = upregulated; green = downregulated. (B) Pathway enrichment scores using MetaboAnalyst 5.0 with 20 input metabolites. These metabolites were significantly elevated in AsPC-1 SERPINB3-high cells. (C) Prediction of upstream regulators for the metabolism in AsPC-1 SERPINB3-high cells. The upstream analysis by IPA using the 20 input metabolites predicts MYC signaling as being involved as a regulator of metabolism in these cells. (D) Heatmap summarizing a metabolome analysis of 116 cancer-related metabolites in SERPINB3-overexpressing MIA PaCa-2 cells, comparing their levels with vector control cells. 36 metabolites were significantly elevated and 5 were decreased (p < 0.05). (E) Pathway enrichment scores using MetaboAnalyst 5.0 with 36 input metabolites. These metabolites were significantly elevated in MIA PaCa-2 SERPINB3-high cells. (F) Prediction of upstream regulators for the metabolism in MIA PaCa-2 SERPINB3-high cells. The upstream analysis by IPA using the 36 input metabolites predicts MYC signaling as being involved as a regulator of metabolism in these cells.

**Supplementary Figure 4.**
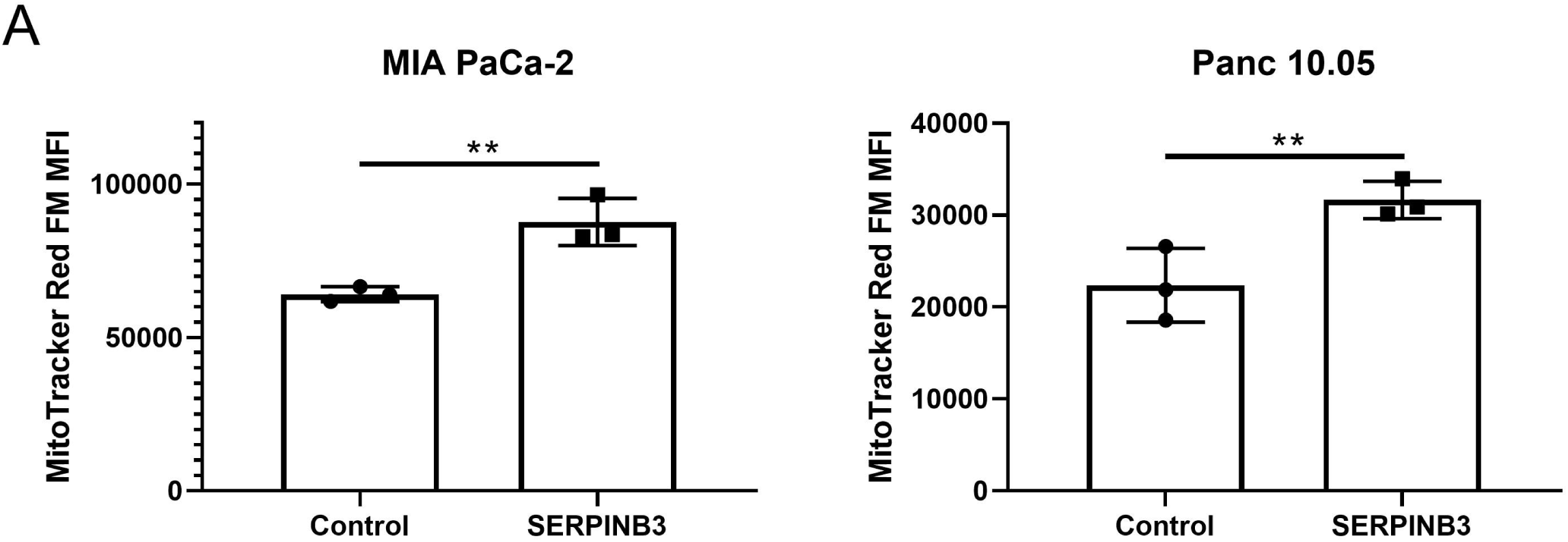
SERPINB3-overexpressing PDAC cells acquire an upregulated mitochondrial membrane potential. Increased mitochondrial membrane potential in SERPINB3-overexpressing PDAC cells. The mitochondrial membrane potential was quantified in cells using MitoTracker Red FM and flow cytometry. Data represent mean ± SD with t-test. *p < 0.05, **p < 0.01, ***p < 0.005.

**Supplementary Figure 5.**
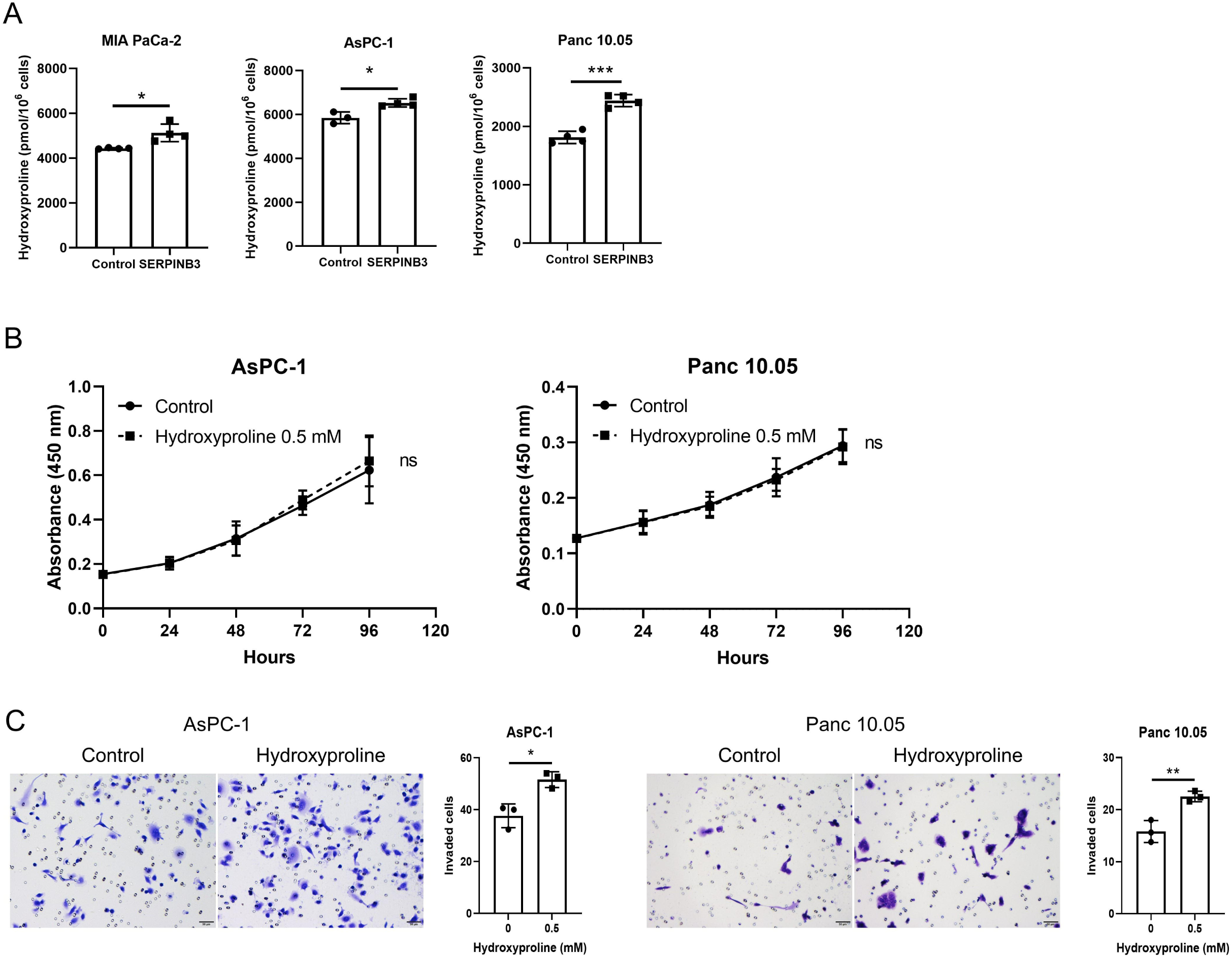
Hydroxyproline increases invasion of PDAC cells. (A) Increased hydroxyproline level in SERPINB3-overexpressing PDAC cells. (B) Hydroxyproline added to the culture medium has no effect on proliferation (CCK-8/WST-8 assay). (C) Hydroxyproline increases invasiveness of PDAC cell lines. Cells that passed through a cell culture insert membrane coated with Matrigel were fixed after 48 hours and counted. Data represent mean ± SD with t-test. *p < 0.05, **p < 0.01, ***p < 0.005.

## References

1. Siegel RL, Miller KD, Wagle NS, Jemal A. Cancer statistics, 2023. CA: A Cancer Journal for Clinicians 2023;73(1):17-48 doi https://doi.org/10.3322/caac.21763.

2. Sung H, Ferlay J, Siegel RL, Laversanne M, Soerjomataram I, Jemal A, et al. Global Cancer Statistics 2020: GLOBOCAN Estimates of Incidence and Mortality Worldwide for 36 Cancers in 185 Countries. CA: A Cancer Journal for Clinicians 2021;71(3):209–49 doi https://doi.org/10.3322/caac.21660.

3. Bailey P, Chang DK, Nones K, Johns AL, Patch A-M, Gingras M-C, et al. Genomic analyses identify molecular subtypes of pancreatic cancer. Nature 2016;531(7592):47–52 doi 10.1038/nature16965.

4. Moffitt RA, Marayati R, Flate EL, Volmar KE, Loeza SGH, Hoadley KA, et al. Virtual microdissection identifies distinct tumor- and stroma-specific subtypes of pancreatic ductal adenocarcinoma. Nature Genetics 2015;47(10):1168–78 doi 10.1038/ng.3398.

5. Collisson EA, Sadanandam A, Olson P, Gibb WJ, Truitt M, Gu S, et al. Subtypes of pancreatic ductal adenocarcinoma and their differing responses to therapy. Nat Med 2011;17(4):500–3 doi 10.1038/nm.2344.

6. Integrated Genomic Characterization of Pancreatic Ductal Adenocarcinoma. Cancer Cell 2017;32(2):185–203.e13 doi 10.1016/j.ccell.2017.07.007.

7. Collisson EA, Bailey P, Chang DK, Biankin AV. Molecular subtypes of pancreatic cancer. Nature Reviews Gastroenterology & Hepatology 2019;16(4):207–20 doi 10.1038/s41575-019-0109-y.

8. Encarnación-Rosado J, Kimmelman AC. Harnessing metabolic dependencies in pancreatic cancers. Nature Reviews Gastroenterology & Hepatology 2021;18(7):482–92 doi 10.1038/s41575-021-00431-7.

9. Ohara Y, Valenzuela P, Hussain SP. The interactive role of inflammatory mediators and metabolic reprogramming in pancreatic cancer. Trends in Cancer 2022;8(7):556–69 doi https://doi.org/10.1016/j.trecan.2022.03.004.

10. Cairns RA, Harris IS, Mak TW. Regulation of cancer cell metabolism. Nature Reviews Cancer 2011;11(2):85–95 doi 10.1038/nrc2981.

11. Dong Y, Tu R, Liu H, Qing G. Regulation of cancer cell metabolism: oncogenic MYC in the driver’s seat. Signal Transduction and Targeted Therapy 2020;5(1):124 doi 10.1038/s41392-020-00235-2.

12. Daemen A, Peterson D, Sahu N, McCord R, Du X, Liu B, et al. Metabolite profiling stratifies pancreatic ductal adenocarcinomas into subtypes with distinct sensitivities to metabolic inhibitors. Proc Natl Acad Sci U S A 2015;112(32):E4410–7 doi 10.1073/pnas.1501605112.

13. Brunton H, Caligiuri G, Cunningham R, Upstill-Goddard R, Bailey U-M, Garner IM, et al. HNF4A and GATA6 Loss Reveals Therapeutically Actionable Subtypes in Pancreatic Cancer. Cell Reports 2020;31(6):107625 doi https://doi.org/10.1016/j.celrep.2020.107625.

14. Espiau-Romera P, Courtois S, Parejo-Alonso B, Sancho P. Molecular and Metabolic Subtypes Correspondence for Pancreatic Ductal Adenocarcinoma Classification. J Clin Med 2020;9(12) doi 10.3390/jcm9124128.

15. Mehla K, Singh PK. Metabolic Subtyping for Novel Personalized Therapies Against Pancreatic Cancer. Clinical Cancer Research 2020;26(1):6–8 doi 10.1158/1078-0432.Ccr-19-2926.

16. Yang S, Tang W, Azizian A, Gaedcke J, Ströbel P, Wang L, et al. Dysregulation of HNF1B/Clusterin Axis Enhances Disease Progression in a Highly Aggressive Subset of Pancreatic Cancer Patients. Carcinogenesis 2022 doi 10.1093/carcin/bgac092.

17. Evans AM, DeHaven CD, Barrett T, Mitchell M, Milgram E. Integrated, Nontargeted Ultrahigh Performance Liquid Chromatography/Electrospray Ionization Tandem Mass Spectrometry Platform for the Identification and Relative Quantification of the Small-Molecule Complement of Biological Systems. Analytical Chemistry 2009;81(16):6656–67 doi 10.1021/ac901536h.

18. Budhu A, Roessler S, Zhao X, Yu Z, Forgues M, Ji J, et al. Integrated Metabolite and Gene Expression Profiles Identify Lipid Biomarkers Associated With Progression of Hepatocellular Carcinoma and Patient Outcomes. Gastroenterology 2013;144(5):1066–75.e1 doi https://doi.org/10.1053/j.gastro.2013.01.054.

19. Zhang G, He P, Tan H, Budhu A, Gaedcke J, Ghadimi BM, et al. Integration of metabolomics and transcriptomics revealed a fatty acid network exerting growth inhibitory effects in human pancreatic cancer. Clin Cancer Res 2013;19(18):4983–93 doi 10.1158/1078-0432.Ccr-13-0209.

20. Wang L, Tang W, Yang S, He P, Wang J, Gaedcke J, et al. NO•/RUNX3/kynurenine metabolic signaling enhances disease aggressiveness in pancreatic cancer. International Journal of Cancer 2020;146(11):3160–9 doi https://doi.org/10.1002/ijc.32733.

21. Mishra P, Tang W, Putluri V, Dorsey TH, Jin F, Wang F, et al. ADHFE1 is a breast cancer oncogene and induces metabolic reprogramming. J Clin Invest 2018;128(1):323–40 doi 10.1172/jci93815.

22. Glynn SA, Boersma BJ, Dorsey TH, Yi M, Yfantis HG, Ridnour LA, et al. Increased NOS2 predicts poor survival in estrogen receptor-negative breast cancer patients. J Clin Invest 2010;120(11):3843–54 doi 10.1172/jci42059.

23. Funamizu N, Hu C, Lacy C, Schetter A, Zhang G, He P, et al. Macrophage migration inhibitory factor induces epithelial to mesenchymal transition, enhances tumor aggressiveness and predicts clinical outcome in resected pancreatic ductal adenocarcinoma. International Journal of Cancer 2013;132(4):785–94 doi https://doi.org/10.1002/ijc.27736.

24. Kato H, Torigoe T. Radioimmunoassay for tumor antigen of human cervical squamous cell carcinoma. Cancer 1977;40(4):1621–8 doi https://doi.org/10.1002/1097-0142(197710)40:4<1621::AID-CNCR2820400435>3.0.CO;2-I.

25. Kato H. Expression and function of squamous cell carcinoma antigen. Anticancer Res 1996;16(4b):2149–53.

26. Petty RD, Kerr KM, Murray GI, Nicolson MC, Rooney PH, Bissett D, et al. Tumor transcriptome reveals the predictive and prognostic impact of lysosomal protease inhibitors in non-small-cell lung cancer. J Clin Oncol 2006;24(11):1729–44 doi 10.1200/jco.2005.03.3399.

27. Vidalino L, Doria A, Quarta S, Zen M, Gatta A, Pontisso P. SERPINB3, apoptosis and autoimmunity. Autoimmun Rev 2009;9(2):108–12 doi 10.1016/j.autrev.2009.03.011.

28. Cannito S, Foglia B, Villano G, Turato C, Delgado TC, Morello E, et al. SerpinB3 Differently Up-Regulates Hypoxia Inducible Factors −1α and −2α in Hepatocellular Carcinoma: Mechanisms Revealing Novel Potential Therapeutic Targets. Cancers 2019;11(12):1933.

29. Cannito S, Turato C, Paternostro C, Biasiolo A, Colombatto S, Cambieri I, et al. Hypoxia up-regulates SERPINB3 through HIF-2α in human liver cancer cells. Oncotarget 2015;6(4):2206–21 doi 10.18632/oncotarget.2943.

30. Turato C, Buendia MA, Fabre M, Redon MJ, Branchereau S, Quarta S, et al. Over-expression of SERPINB3 in hepatoblastoma: A possible insight into the genesis of this tumour? European Journal of Cancer 2012;48(8):1219–26 doi https://doi.org/10.1016/j.ejca.2011.06.004.

31. Turato C, Cannito S, Simonato D, Villano G, Morello E, Terrin L, et al. SerpinB3 and Yap Interplay Increases Myc Oncogenic Activity. Scientific Reports 2015;5(1):17701 doi 10.1038/srep17701.

32. Correnti M, Cappon A, Pastore M, Piombanti B, Lori G, Oliveira DVPN, et al. The protease-inhibitor SerpinB3 as a critical modulator of the stem-like subset in human cholangiocarcinoma. Liver International 2022;42(1):233–48 doi https://doi.org/10.1111/liv.15049.

33. Lauko A, Volovetz J, Turaga SM, Bayik D, Silver DJ, Mitchell K, et al. SerpinB3 drives cancer stem cell survival in glioblastoma. Cell Rep 2022;40(11):111348 doi 10.1016/j.celrep.2022.111348.

34. Pontisso P. Role of SERPINB3 in hepatocellular carcinoma. Ann Hepatol 2014;13(6):722–7.

35. Sheshadri N, Catanzaro JM, Bott AJ, Sun Y, Ullman E, Chen EI, et al. SCCA1/SERPINB3 Promotes Oncogenesis and Epithelial–Mesenchymal Transition via the Unfolded Protein Response and IL6 Signaling. Cancer Research 2014;74(21):6318–29 doi 10.1158/0008-5472.Can-14-0798.

36. Catanzaro JM, Sheshadri N, Pan J-A, Sun Y, Shi C, Li J, et al. Oncogenic Ras induces inflammatory cytokine production by upregulating the squamous cell carcinoma antigens SerpinB3/B4. Nature Communications 2014;5(1):3729 doi 10.1038/ncomms4729.

37. Chandriani S, Frengen E, Cowling VH, Pendergrass SA, Perou CM, Whitfield ML, et al. A core MYC gene expression signature is prominent in basal-like breast cancer but only partially overlaps the core serum response. PLoS One 2009;4(8):e6693 doi 10.1371/journal.pone.0006693.

38. Terunuma A, Putluri N, Mishra P, Mathé EA, Dorsey TH, Yi M, et al. MYC-driven accumulation of 2-hydroxyglutarate is associated with breast cancer prognosis. J Clin Invest 2014;124(1):398–412 doi 10.1172/jci71180.

39. Smid M, Wang Y, Zhang Y, Sieuwerts AM, Yu J, Klijn JGM, et al. Subtypes of Breast Cancer Show Preferential Site of Relapse. Cancer Research 2008;68(9):3108–14 doi 10.1158/0008-5472.Can-07-5644.

40. Yu K, Chen B, Aran D, Charalel J, Yau C, Wolf DM, et al. Comprehensive transcriptomic analysis of cell lines as models of primary tumors across 22 tumor types. Nature Communications 2019;10(1):3574 doi 10.1038/s41467-019-11415-2.

41. Furuno T, Kanno T, Arita K, Asami M, Utsumi T, Doi Y, et al. Roles of long chain fatty acids and carnitine in mitochondrial membrane permeability transition. Biochem Pharmacol 2001;62(8):1037–46 doi 10.1016/s0006-2952(01)00745-6.

42. Ray CA, Black RA, Kronheim SR, Greenstreet TA, Sleath PR, Salvesen GS, et al. Viral inhibition of inflammation: Cowpox virus encodes an inhibitor of the interleukin-1β converting enzyme. Cell 1992;69(4):597–604 doi https://doi.org/10.1016/0092-8674(92)90223-Y.

43. Schick C, Pemberton PA, Shi G-P, Kamachi Y, Çataltepe S, Bartuski AJ, et al. Cross-Class Inhibition of the Cysteine Proteinases Cathepsins K, L, and S by the Serpin Squamous Cell Carcinoma Antigen 1:□ A Kinetic Analysis. Biochemistry 1998;37(15):5258–66 doi 10.1021/bi972521d.

44. Sun Y, Sheshadri N, Zong WX. SERPINB3 and B4: From biochemistry to biology. Semin Cell Dev Biol 2017;62:170–7 doi 10.1016/j.semcdb.2016.09.005.

45. Collie-Duguid ESR, Sweeney K, Stewart KN, Miller ID, Smyth E, Heys SD. SerpinB3, a new prognostic tool in breast cancer patients treated with neoadjuvant chemotherapy. Breast Cancer Research and Treatment 2012;132(3):807–18 doi 10.1007/s10549-011-1625-9.

46. Ullman E, Pan JA, Zong WX. Squamous cell carcinoma antigen 1 promotes caspase-8-mediated apoptosis in response to endoplasmic reticulum stress while inhibiting necrosis induced by lysosomal injury. Mol Cell Biol 2011;31(14):2902–19 doi 10.1128/mcb.05452-11.

47. Casciano JC, Perry C, Cohen-Nowak AJ, Miller KD, Vande Voorde J, Zhang Q, et al. MYC regulates fatty acid metabolism through a multigenic program in claudin-low triple negative breast cancer. British Journal of Cancer 2020;122(6):868–84 doi 10.1038/s41416-019-0711-3.

48. Pacilli A, Calienni M, Margarucci S, D’Apolito M, Petillo O, Rocchi L, et al. Carnitine-Acyltransferase System Inhibition, Cancer Cell Death, and Prevention of Myc-Induced Lymphomagenesis. JNCI: Journal of the National Cancer Institute 2013;105(7):489–98 doi 10.1093/jnci/djt030.

49. Melone MAB, Valentino A, Margarucci S, Galderisi U, Giordano A, Peluso G. The carnitine system and cancer metabolic plasticity. Cell Death & Disease 2018;9(2):228 doi 10.1038/s41419-018-0313-7.

50. McCann MR, George De la Rosa MV, Rosania GR, Stringer KA. L-Carnitine and Acylcarnitines: Mitochondrial Biomarkers for Precision Medicine. Metabolites. Volume 112021.

51. Li S, Gao D, Jiang Y. Function, Detection and Alteration of Acylcarnitine Metabolism in Hepatocellular Carcinoma. Metabolites. Volume 92019.

52. Muoio Deborah M, Noland Robert C, Kovalik J-P, Seiler Sarah E, Davies Michael N, DeBalsi Karen L, et al. Muscle-Specific Deletion of Carnitine Acetyltransferase Compromises Glucose Tolerance and Metabolic Flexibility. Cell Metabolism 2012;15(5):764–77 doi https://doi.org/10.1016/j.cmet.2012.04.005.

53. Li J-L, Wang Q-Y, Luan H-Y, Kang Z-C, Wang C-B. Effects of L-carnitine against oxidative stress in human hepatocytes: involvement of peroxisome proliferator-activated receptor alpha. Journal of Biomedical Science 2012;19(1):32 doi 10.1186/1423-0127-19-32.

54. Fink MA, Paland H, Herzog S, Grube M, Vogelgesang S, Weitmann K, et al. L-Carnitine–Mediated Tumor Cell Protection and Poor Patient Survival Associated with OCTN2 Overexpression in Glioblastoma Multiforme. Clinical Cancer Research 2019;25(9):2874–86 doi 10.1158/1078-0432.Ccr-18-2380.

55. Altun ZS, Güneş D, Aktaş S, Erbayrktar Z, Olgun N. Protective Effects of Acetyl-l-Carnitine on Cisplatin Cytotoxicity and Oxidative Stress in Neuroblastoma. Neurochemical Research 2010;35(3):437–43 doi 10.1007/s11064-009-0076-8.

